# Ecological constraints and evolutionary trade-offs shape nitrogen fixation across habitats

**DOI:** 10.1101/2025.05.20.655134

**Authors:** Morgan S. Sobol, Aya S. Klos, Cécile Ané, Katherine D. McMahon, Betül Kaçar

## Abstract

From its earliest beginnings, life’s expansion into new habitats has been profoundly shaped by its reciprocal interactions with Earth’s changing environments. Understanding how ancient metabolisms co-evolved with their environments requires uncovering the ecological and evolutionary processes that structured the functionally linked genes and networks underlying these metabolisms. Here, we focus on nitrogen (N_2_) fixation, one of life’s most critical metabolisms, and investigate the drivers of complexity in its associated gene machinery today. We used a large-scale comparative genomics framework to construct a comprehensive catalog of extant N_2_ fixation-associated genes and assessed their distribution across diverse microbial genomes and environmental backgrounds. Genomes enriched in N_2_ fixation genes generally have larger genome sizes, broader metabolic capabilities, wider habitat ranges, and are predominantly associated with mesophilic and aerobic lifestyles. Evolutionary reconstructions reveal a pattern of early gene gains in ancestral diazotrophs followed by lineage-specific gene losses in later diverging taxa, suggesting evolutionary trade-offs shaped by changing environments. These findings demonstrate that the evolution of N_2_ fixation has been intertwined with the composition and organization of the genes supporting the overarching N_2_ metabolism, driven by feedback between genome evolution and shifting environmental and ecological conditions.

## Introduction

Biological nitrogen fixation (“N_2_ fixation”) is an ancient metabolism characterized by the reduction of atmospheric N_2_ to bioavailable ammonia [1, 2]. Nitrogenases are the only known enzymes capable of catalyzing this energy-intensive reaction, giving them a crucial historical role expanding the biosphere through nitrogen supply [3, 4].

Phylogenetic reconstructions support a shared ancestry among five major clades of nitrogenases that have diversified over time [5]. These include three isozymes defined by the metal in their active site cofactors, molybdenum (Mo), vanadium (V), or iron (Fe), and the three Mo-nitrogenase clades that differ in host ecology and taxonomy [2, 6, 7]. Structural variations distinguish the Mo-nitrogenase clades and align with historic transitions in environmental oxygen levels [6]. Despite this diversification, nitrogenase’s core structure [6] and catalytic mechanism [8, 9] have remained largely conserved over ∼3 billion years of evolution [10, 11].

Outside the main nitrogenase complex (*nifHDK* for Mo-nitrogenases), N_2_ fixation depends on a myriad of other proteins that have coevolved with it throughout the history of this metabolism [12, 13]. These include proteins involved in cofactor assembly (*nifBEN*) [14], electron transfer (*nifFJ*), transcriptional regulation (*nifALI_1_I_2_*), and biosynthesis (*nifXQUVYS*) [15, 16]. Beyond the *nif* cluster, numerous other gene families contribute to N_2_ fixation, including those involved in oxygen protection (*nod, ahp, coo, cow, sod*), electron donation and transfer (*fix, rng, fdx*), transcriptional regulation (*draGT, fixJLK, regRS, ntc, ntr*), metal transport (*mod, feo, cys*), iron-sulfur cluster assembly (*suf, isc, fdx*) [17], as well as nitrogen metabolism (*gln, nir, nap, hox*) [18] (**Figure 1**). Additional genes with potential roles in N_2_ fixation remain functionally uncharacterized [19].

**Figure 1.**
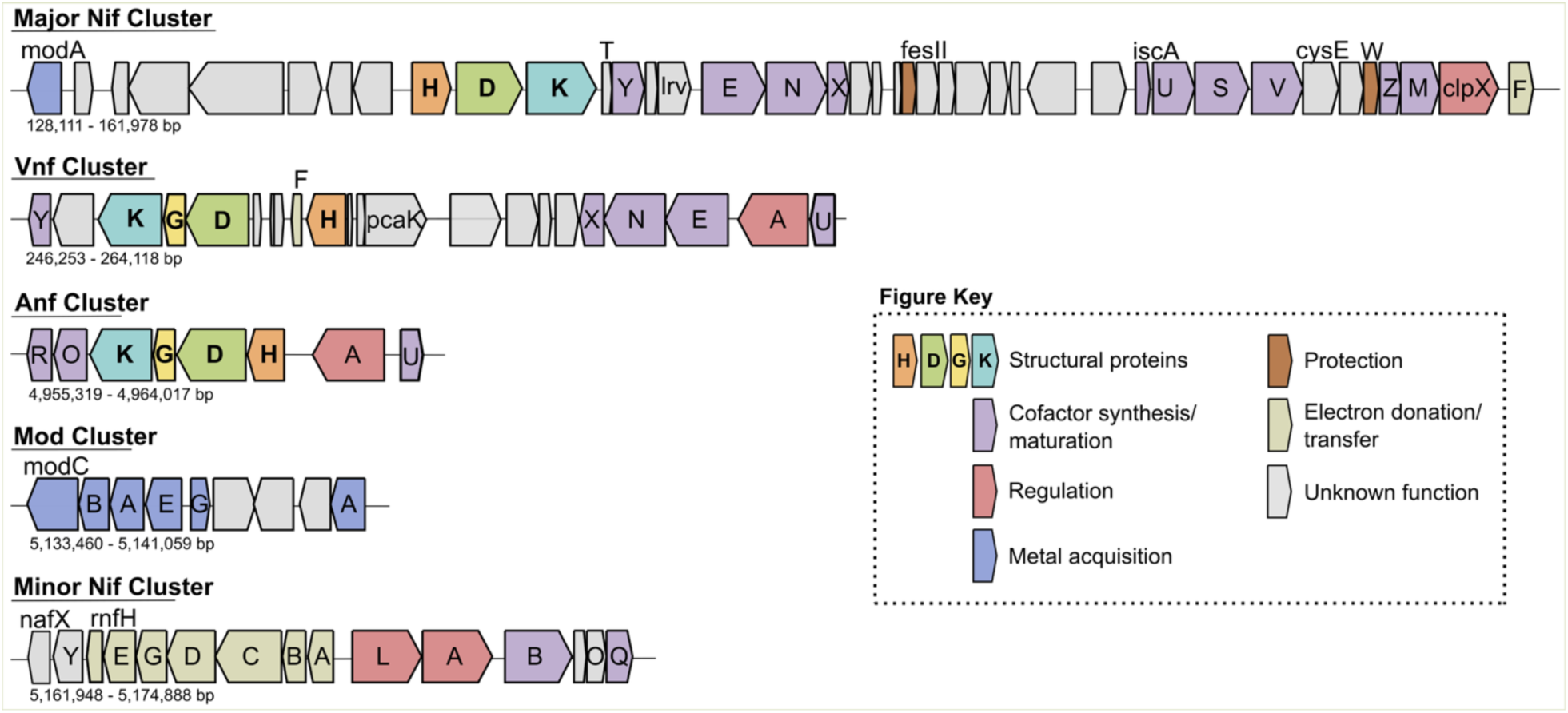
Structure of the major and minor molybdenum nitrogenase (Nif) clusters, vanadium nitrogenase (Vnf) and iron nitrogenase (Anf) clusters, and the molybdenum acquisition (Mod) cluster in model diazotroph *Azotobacter vinelandii* DJ. Genome positions and gene sizes for each cluster are based on KEGG genome reference T00885 for *A*. *vinelandii* DJ.

The composition of the complex genetic machinery responsible for N_2_-fixation varies widely among diazotrophs, with the simplest gene sets found in hyperthermophilic methanogens and the most elaborate networks in aerobes [12]. Previous studies have highlighted the diverse evolutionary trajectories of nitrogenase-associated genes [13] and revealed a dynamic history of gene acquisitions and losses linked to the emergence of aerobic metabolisms [12]. Nevertheless, a central challenge remains to reconstruct the evolutionary history of N_2_ fixation in deep time and relate it to the ecological diversification of diazotrophs. Here, we address this challenge using large-scale comparative genomics and show how the evolution of N_2_ fixation genes enabled diazotrophs to expand into diverse and increasingly oxygenated environments.

## Methods

### Genome selection

The reference phylogeny Web of Life (WoL) [20] was selected as the basis of our study because of its extensive genome coverage and phylogenetic reconstruction. For our analysis, we kept WoL genomes with ≥90% completeness and <5% contamination, following quality thresholds established by [21]. Protein fasta files associated with the genome accessions (named “*_protein.faa.gz”) were downloaded from NCBI with eutils [22] (accessed July 17th, 2023). Genomes missing available NCBI protein fasta files were excluded from analysis. After filtering for high-quality genomes and removing those without annotated fasta files, the WoL genome dataset was reduced from 10,575 genomes to 7,458. Nucleotide sequences of noncoding RNA genes were downloaded if they were available at GenBank or RefSeq and the 16S rRNA genes were extracted.

### Nitrogen fixation gene catalog curation

Protein fasta files were functionally annotated using the Kyoto Encyclopedia of Genes and Genomes (KEGG) database [23]. Specifically, the translated amino acid sequences were annotated with KEGG orthology (KO) identifiers by profiling each genome against KEGG hidden Markov models (KEGG release 106) for all KOs [24] (cutoff E-value = 1e- 5) using *hmmscan* in HMMER version 3.2.2 (http://hmmer.org/). Custom Python scripts were used to parse through each genome’s *hmmscan* output and generate the final KO presence-absence matrix. The presence/absence matrix was subset to genes known to be involved in N_2_ fixation (and which had KO identifiers), which were collected through an extensive literature search that included foundational studies with thorough analyses of genes involved in N_2_ fixation (**Supplemental Table 1**). Gene involvement was supported by functional confirmation, for example by evidence from protein interactions with nitrogenase, gene knockout phenotypes, or differential gene expression under N_2_ fixation.

### Species tree construction and visualization

The WoL species tree was subsetted to genomes annotated with *nifHDK* using the *drop.tip* function from the APE package [25] in R. We additionally confirmed the presence of nitrogenase by phylogenetic placement of concatenated *nifDK* sequences, as described in reference [5], from each genome. The resulting species tree, the gene presence-absence matrix, and trait data were plotted using the ggTree R package (version 3.6.2) [26].

### Trait data curation

Optimal growth temperature (OGT), oxygen, pH preference, and habitat preference trait data were collected for each genome by matching the GenBank accession number to BacDive [27], NCBI BioProject [28], or GOLD JGI [29] as well as matching the NCBI taxon IDs to the Madin et al (2020) phenotypic trait database [30]. Trait data was also predicted for all genomes. OGTs were predicted using the OGT_prediction tool [31]. The regression model used included both Bacteria and Archaea, but excluded using information of the 16S rRNA gene and genome size (superkingdom-all_species-all_features_ex_genome_size_and_rRNA.txt). The *Oxyphen* package was used for oxygen metabolism prediction based on the number of oxygen utilizing genes in each genome [32]. Predictions for pH were done with the Ramoneda et al. (2023) prediction model which is based on genes that influence pH tolerance [33]. Habitat preference and coverage were determined with ProkAtlas [34] by mapping the 16S rRNA gene sequences to the ProkAtlas metagenome database. Barrnap version 0.9 [35] was used to extract 16S rRNA sequences for genomes with missing annotations. Using a custom python script, the longest 16S rRNA sequence was extracted for each genome as its representative 16S rRNA gene sequence to correct for errors in sequence length due to sequencing and assembly bias. Twenty-one genomes without a 16S rRNA gene sequence were excluded. A nucleotide identity of 97% and a query length match of at least 150 base pairs were used as thresholds. Habitat preference, categorized into freshwater, saline, terrestrial, plant-associated, non-plant host-associated, and hydrothermal, was defined as the environment with the highest 16S abundance for each genome. Habitat coverage was determined by summing up the total number of habitats where a genome was considered present and dividing it by the total number of habitats (n=100) investigated by ProkAtlas. The presence of KEGG modules was predicted using the KEGG KO annotations with MetaPathPredict (version 0) [36]. Metabolic coverage was calculated as the total number of KEGG modules present in a genome divided by the total number of modules (n=194) searched for with MetaPathPredict. Data collected from databases and predicted data were cross-referenced and manually annotated to provide the final trait data for each genome.

### Data analysis and statistics

We compared the mean number of N_2_ fixation genes across specific traits (oxygen, temperature, and habitat preferences) with non-parametric methods using the *stat_compare_means* function in the ggpubr R package (version 0.6.0) [37]. For the categorical traits habitat, oxygen, and temperature preference, we first applied a Kruskal-Wallis test, followed by pairwise Wilcoxon rank-sum tests with Benjamini-Hochberg correction. For temperature preference (categories: mesophilic and thermophilic) we used the Wilcoxon rank-sum test directly. Compact letter display was used to report pairwise groups whose means are discernibly different, based on a *P*-value of 0.05.

The gene presence-absence matrix was used in phylogenetic principal-component analysis (PCA) performed with the *phyl.pca* function (method = “BM”, mode =”corr”) in phytools R package [38]. This was done to visualize similarities and dissimilarities between genomes regarding N_2_ fixation gene content while accounting for phylogenetic relationships. This approach reduces the risk of over-interpreting similarities simple due to common ancestry among taxa. The first two components were retained for visualization. Pearson correlation between the presence-absence of N_2_ fixation genes was calculated with the *cor* function in the stats R package [39] and plotted using corrplot [40] with hierarchical clustering and default *P*-value of 0.05, to identify patterns of gene co-occurrence.

### Evolutionary trends analysis

In order to test whether certain traits shared a coevolutionary history with N_2_ fixation gene abundance, phylogenetic linear regression models were fit using the *phylolm* function in the phylolm R package (version 2.62) [41], using Pagel’s lambda model of phylogenetic correlation. The lambda parameter (λ) ranges from 0 (no phylogenetic signal) to 1 (high phylogenetic signal) in the response trait when the model has no covariates, or in the residuals when covariates are included. When assessing phylogenetic signal within individual clades, those represented by fewer than 20 genomes were not included in the analysis, as statistical power to detect signal decreases substantially in trees with fewer than 20 branches [42, 43]. Non-phylogenetic linear regressions were performed using the *lm* function in the stats R package [39]. The *distRoot* function in the adephylo R package [44] was used to calculate the root-to-tip patristic distances by measuring the total length of branches (in terms of the number of substitutions per site) from the root of the species tree to each tip.

Gene gain/loss was determined with Count [45] using a non-binary count matrix [46] for all accessory (non-structural) *nif* genes. Wagner parsimony with a gain-to-loss penalty ratio of 2 was applied to infer the most parsimonious scenario of gene presence at internal nodes, as described previously [47]. This ratio is widely used because recurring observations indicate that gene loss occurs at least twice as frequently than gene gain [48–50]. Ancestral trait reconstructions were performed on an ultrametric and dichotomous tree, which was made using the *multi2di* and *chronos* (model = “relaxed”) functions of the APE package [25] in R. The *fitdiscrete* function in geiger [51] was used for discrete reconstructions of oxygen preference, which were mapped onto the phylogeny with the *simmap* function in phytools [38]. The *fastAnc* function in phytools was used for continuous variable reconstructions of metabolic and habitat diversity, which was mapped onto the tree using the *contMap* function in phytools [38].

### Sampling genomes from non-diazotrophs

To assess whether the observed co-evolutionary trends between traits and N_2_ fixation gene abundance were specific to diazotrophs, we analyzed a random sample of non-diazotrophic genomes using the same phylogenetic linear regression methodology described above. Among the 6,785 genomes missing one or more of the catalytic *nifHDK* subunits, 700 were randomly selected with the base R *sample* function (version 4.4.2) [39], for further analysis (**Supplemental Table 9**). Trait data (including OGT, oxygen status, pH preference, and habitat preference) and gene presence/absence for N_2_ fixation-related genes were collected as described for diazotrophs (**Supplemental Tables 9, 10**). Fifty-one genomes without 16S rRNA gene sequences were excluded from analyses involving habitat coverage.

## Results

### Distribution of nitrogenases across the tree of life

We used a large prokaryote reference phylogeny, “WoL” [20], containing over 7,000 high- quality genomes [21] to first assess the representative diversity and distribution of nitrogenase today. A total of 673 genomes (∼9.0%) were found to contain nitrogenase (**Supplemental Table 2**) based on the presence of the catalytic HDK subunits of nitrogenase (**Figure 2A**). Our estimate of nitrogenase diversity falls within the range of recent estimates (∼5.9% to 16%) [7, 12, 52, 53]. Here, approximately 46.9% of these genomes belonged to Proteobacteria (in order of abundance: Alpha>Delta>Gamma>Beta>Epsilon>Acidithiobacillia), while the remaining top three phyla included Firmicutes (17.3%), Cyanobacteria (12.6%), and Euryarchaeota (11.4%). Amongst the nitrogenase-containing genomes, molybdenum (Mo)-only nitrogenase made up the majority (90.3%), followed by genomes with both Mo-nitrogenase and iron (Fe) nitrogenase (5.9%), genomes with both Mo-nitrogenase and vanadium (V) nitrogenase (2.2%), and lastly with 1.5% of genomes containing all three nitrogenases (**Figure 1**) (**Figure 2B**). Genomes with V-nitrogenase were found in 6 phyla: Euryarchaeota, Firmicutes, Alphaproteobacteria, Deltaproteobacteria, Cyanobacteria, and Gammaproteobacteria. Fe-nitrogenases were found in 11 phyla: Firmicutes, Epsilonproteobacteria, Alphaproteobacteria, Gammaproteobacteria, Verrucomicrobia, Euryarchaeota, Deltaproteobacteria, Gammaproteobacteria, Bacteroidetes, Spirochaetes, and Chlorobi. Euryarchaeota represented 50% of the genomes with all three nitrogenases; the remaining phyla included Alphaproteobacteria, Firmicutes, and Gammaproteobacteria.

**Figure 2.**
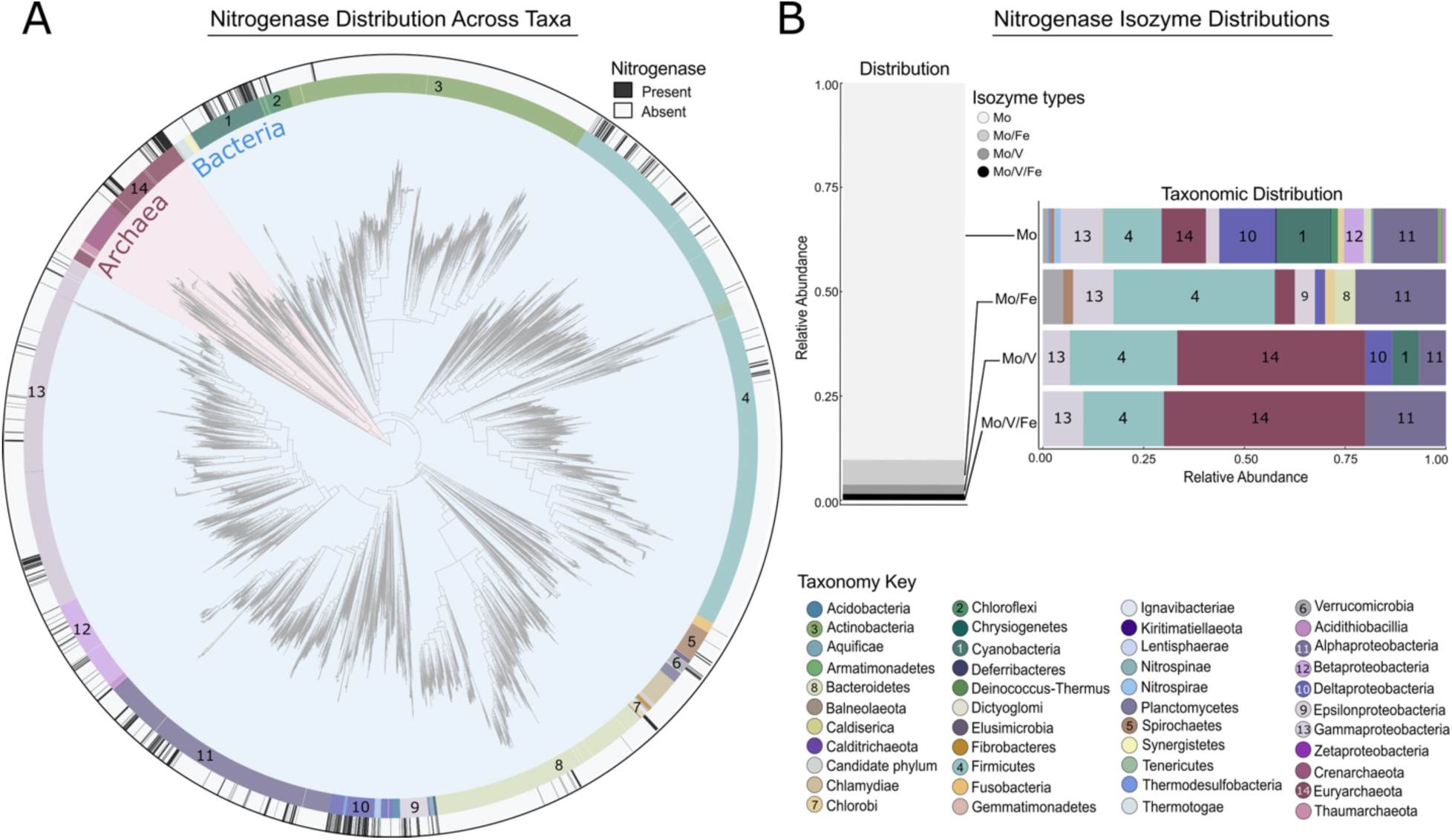
Diversity and phylogenetic distribution of nitrogenase. **A)** The overall distribution of nitrogenase across the microbial diversity from the Zhu et al (2019) Web of Life (WoL) phylogeny [20]. A total of 7,458 high-quality genomes were assessed. Of these, ∼9.0% of all genomes (n=673) encoded genes for the full nitrogenase complex (*nifHDK*). Black bars on the outer ring represent the presence of nitrogenase. Numbers reflect the top 14 taxonomic groups in the dataset with the highest proportion of genomes with nitrogenase. The phylogeny was plotted with iTOL [54]. **B)** Proportions and taxonomic distributions of the different nitrogenase isozymes across WoL. Mo-nitrogenase constitutes the majority of nitrogenases identified, followed by Fe-nitrogenase, and then V-nitrogenase in abundance. Only ∼1% of genomes contained all three nitrogenases, but ∼50% of genomes with all three isozymes belonged to Euryarchaeota.

### An N_2_ fixation genomic and gene-centric ecological catalog

We screened all 673 genomes with the full *nifHDK* complement for 93 known genes (“machinery”) involved in N_2_ fixation (**Table 1, Supplemental Table 3, Supplemental Figure 1**). In Figure 3, genes are grouped by functional category in relation to N_2_ fixation, and the relative abundance of genes within each category is shown. Excluding the structural genes *nifHDK*, the total number of N_2_ fixation genes detected per genome ranged from 14 to 67 genes. The largest N_2_ fixation gene repertoires were observed in the Alphaproteobacteria *Rhodopseudomonas palustris CGA009* and the Gammaproteobacteria *Azotobacter vinelandii* DJ, which contained 67 and 65 genes, respectively. In contrast, the archaea taxa *Methanococcus aeolicus* Nankai-3, *Methanothermococcus okinawensis* IH1 and *Methanocaldococcus infernus* ME each possessed the smallest repertoires with 14 genes. Across the species phylogeny, the number of N_2_ fixation genes per genome scaled positively with branch lengths, indicating lineages with greater cumulative evolutionary change tend to possess larger N_2_ fixation gene complements (**Supplemental Figure 2**).

**Figure 3.**
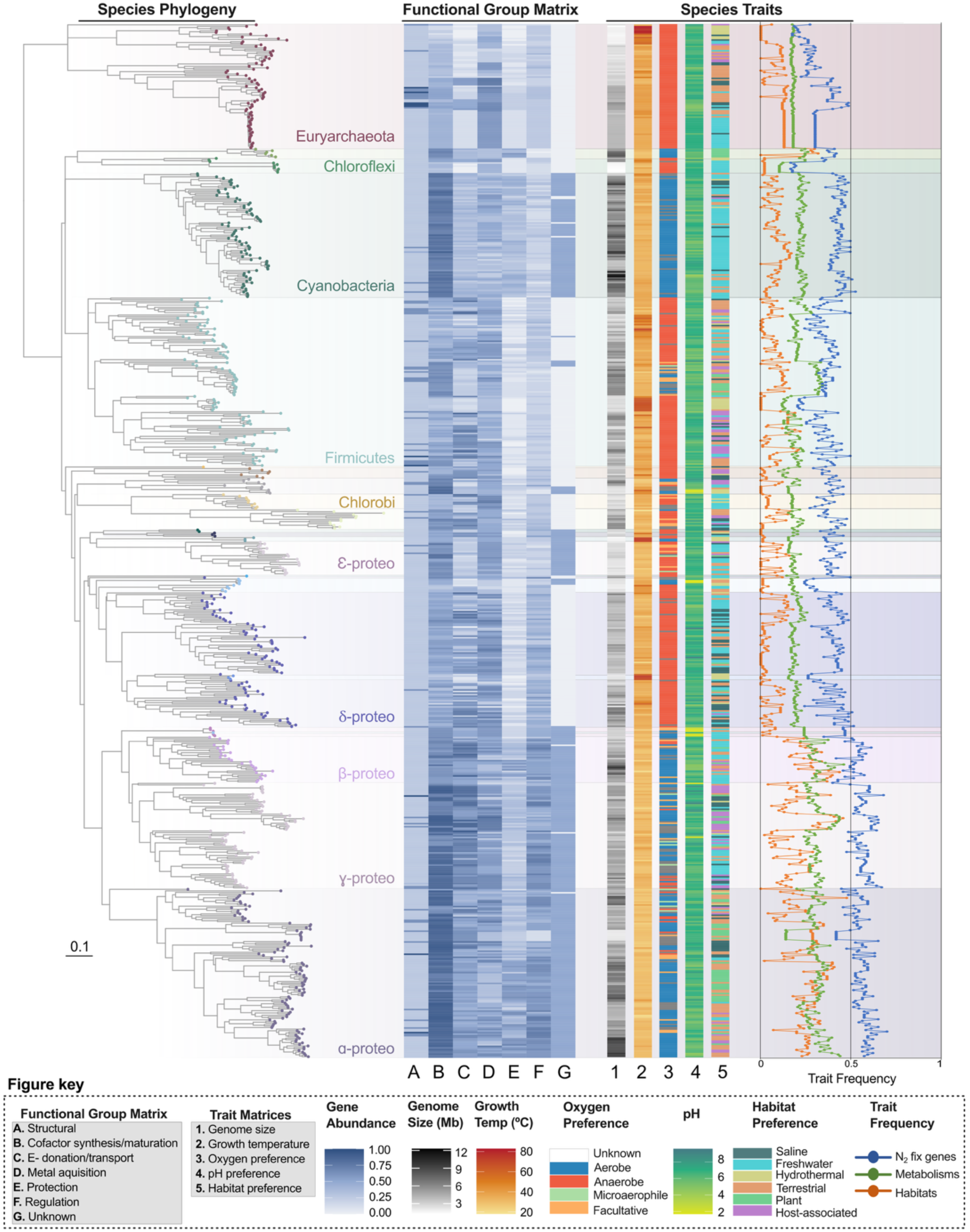
Distribution of N_2_ fixation machinery mapped onto the phylogeny of nitrogenase-containing genomes, highlighting associations with species-associated ecological (habitat preference, genome size, coverage of metabolic pathways) and physiological traits (oxygen, temperature, and pH preferences). N_2_ fixation gene abundance correlates with those that are mesophilic, prefer aerobic and/or facultative lifestyles, possess larger genome sizes, broader metabolic capabilities, and occupy a wider range of habitats. For this figure, N_2_ fixation genes were organized into functional categories and the relative abundance of genes for each category calculated. The full gene matrix visualization can be found in Supplemental Figure 1 and the gene matrix and associated trait categories in Supplemental Table 3. Trait frequencies were calculated as the relative abundances of N_2_ fixation genes (blue line), metabolic pathways (green line), and habitats (orange line) associated with genome hosts. The phylogeny and matrices were plotted using the ggTree R package (version 3.6.2) [26].

**Table 1.**
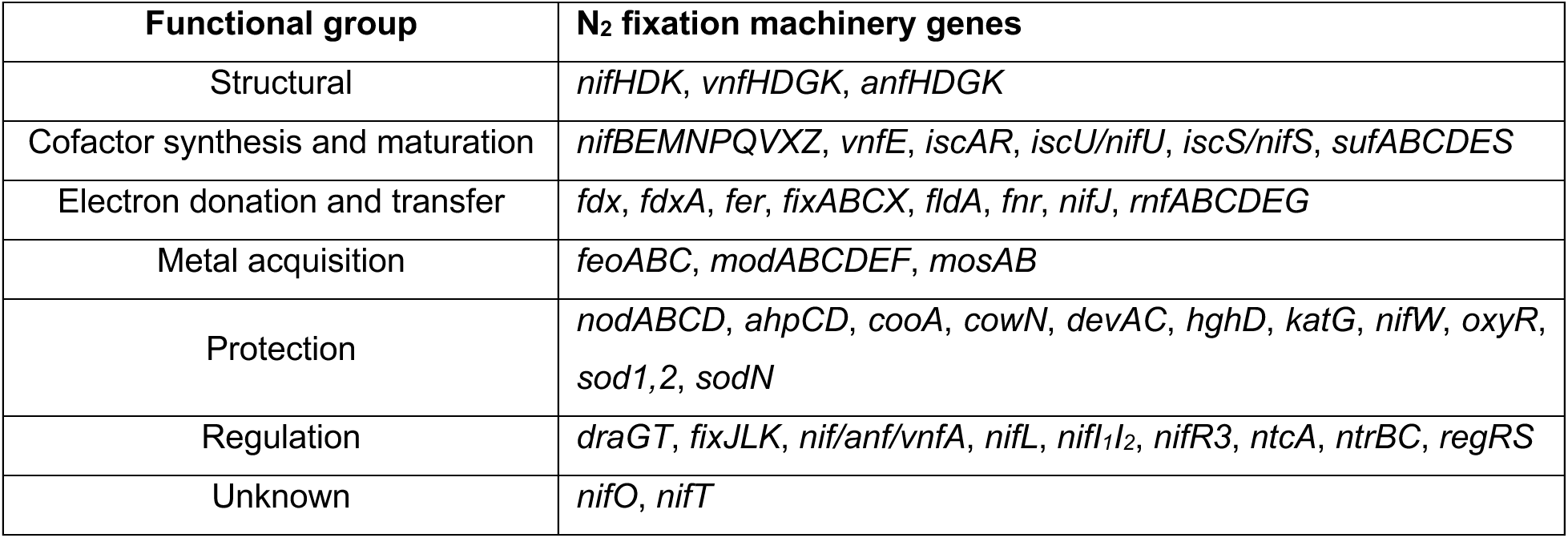
Functional groupings of N_2_ fixation machinery genes.

To explore the physiological and ecological diversity of diazotrophs identified in this study, we mapped their habitat preferences, oxygen tolerance, optimal growth pH and temperature, metabolic versatility, and genome size onto a phylogenetic tree depicting relationships among diazotroph lineages (**Figure 3**).

Amongst nitrogenase-containing genomes, 47.5% belonged anaerobes, 31.6% to aerobes, 5.3% to facultative species, and 1.5% to microaerophiles. The remaining 14% could not be classified due to missing data or uncertainty in predictions. Temperature preferences further define nitrogenase distribution with most genomes (91.2%) classified as mesophiles favoring growth between 18°C and 45°C. The remaining ∼8.8% nitrogenase-containing genomes belonged to thermophiles that grow best at temperatures above 45°C. Similarly, neutrophiles (pH 5.5-8.0) dominated the pH preference dataset (87.2%), whereas acidophiles and alkaliphiles were comparatively rare.

Habitat distributions varied across taxa. Freshwater and terrestrial environments were the most common inferred habitats of preference, encompassing 32.5% and 26.2% of genomes, respectively. Other environments included saline systems (14.3%), non-plant host associations (11.4%), plant-associated niches (8.8%), and hydrothermal systems (6.8%). These patterns likely both reflect genuine ecological preferences and uneven environmental sampling, which may skew our understanding of where nitrogenase-containing organisms are most abundant in nature.

Diazotrophs also differed in their metabolic and ecological breadth. Metabolic coverage ranged from 9.8 to 46.4% with an average of 23.2%, and habitat coverage varied from 0 to 48% with an average of 12%, excluding the ∼3% of genomes for which habitat data were unavailable. Genome size spanned from 1.3 to 12.3 Mbp with a mean of 4.6 Mbps.

### Physiological, metabolic, and habitat preferences vary with N_2_ fixation genetic complexity

Next, we examined how the number of N_2_ fixation genes per genome relates to environmental and physiological traits, to identify the factors influencing the genetic complexity of the N_2_ fixation machinery. Here, genetic complexity is defined as the proportion of N_2_ fixation genes detected within a given genome.

We observed that genomes sharing similar temperature, oxygen, and habitat preferences had comparable numbers of N_2_ fixation genes (**Figure 3**). Specifically, thermophiles and anaerobes contained significantly fewer genes than their mesophilic (p.adj=8.60×10^-23^) and aerobic (p.adj=4.80×10^-45^) counterparts (**Figure 4AB**). Consistently, taxa preferring hydrothermal environments on average have fewer N_2_ fixation genes than those from other habitats (**Figure 4C**). No significant differences were detected among terrestrial and freshwater, saline, or non-plant host-associated environments. Plant-associated taxa exhibited the highest average of N_2_ fixation gene counts, exceeding those from other environments except non-plant host-associated taxa.

**Figure 4.**
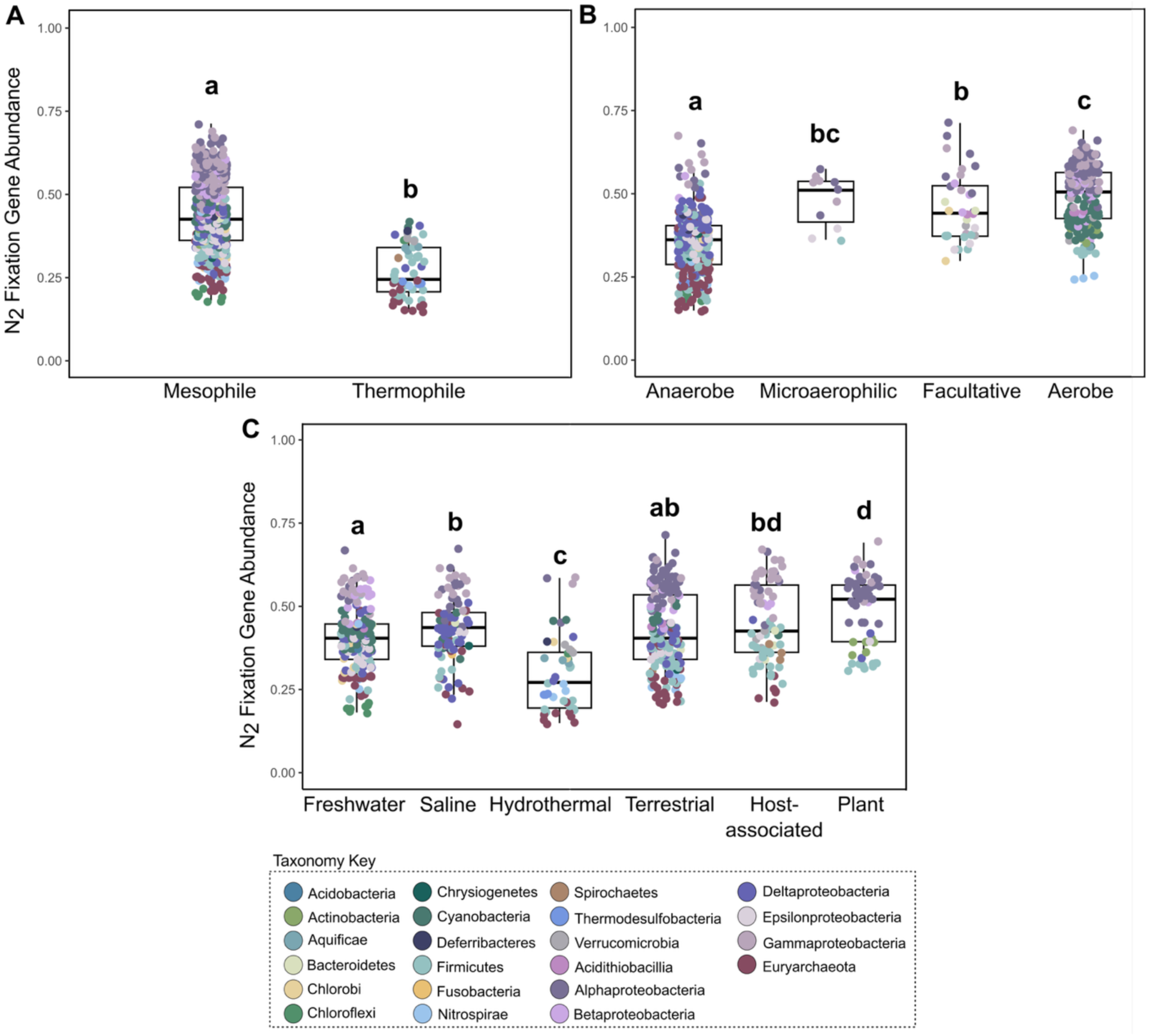
N_2_ fixation gene abundance varies across traits. The number of genes was compared across **A)** temperature**, B)** oxygen, and **C)** habitat preferences. Thermophiles and anaerobes have discernably fewer N_2_ fixation genes than mesophiles and all other oxygen phenotypes, respectively. Genomes associated with hydrothermal environments have on average, the fewest number of N_2_ fixation genes, whereas plant-associated genomes have the highest average. Pairwise differences between means were calculated using non-parametric Wilcoxon rank-sum tests. For each panel, the means of those that do not differ with statistical evidence share the same letter, whereas those that do not share a letter are significantly different (*P*<0.05). Boxplots were plotted with the ggpubr R package [37].

### N_2_ fixation genetic complexity as a predictor of co-evolutionary trait associations

To test the degree to which traits are phylogenetically conserved versus independently evolved, we measured phylogenetic signal for gene abundance, habitat coverage, metabolic coverage, genome size, temperature, pH, and oxygen preference across the species phylogeny using Pagel’s lambda (λ), as implemented in the *phylolm* R package [41]. All traits had a moderate to high phylogenetic signal (λ≥0.69; **Supplemental Table 4**), indicating closely related genomes tend to share more similar trait values than those drawn randomly from the phylogeny.

We further assessed phylogenetic signal within individual phyla to test for lineage-specific evolutionary patterns and differences in selection pressures. This analysis clarified whether the overall phylogenetic signal measured across the species phylogeny was disproportionately driven by particular phyla. Because some phyla had too few genomes (<20), phylum-specific analyses were restricted to Alpha-, Beta-, Delta-, Gammaproteobacteria, as well as Cyanobacteria, Firmicutes, and Euryarchaeota. Genome size, habitat coverage, and metabolic coverage all exhibited strong phylogenetic signal across these major nitrogenase-containing phyla, (λ≥0.8, **Supplemental Table 6**), indicating that these traits are broadly conserved across these lineages, rather than confined to a single clade. N_2_ fixation gene abundance likewise showed a high λ (>0.8) within most phyla with the exception of Cyanobacteria (λ = 0.34), potentially due to limited variability of N_2_ fixation gene sets within this clade.

By contrast, no phylogenetic signal for temperature was detected in Gammaproteobacteria (λ=0.001), consistent with their largely mesophilic lifestyle, whereas other phyla retained strong signals (λ>0.8). Oxygen tolerance showed little phylogenetic structure in either Gammaproteobacteria or Firmicutes. In contrast, pH exhibited strong phylogenetic conservation in Cyanobacteria, Euryarcheaota, and Gamma- and Betaproteobacteria, but low conservation in Deltaproteobacteria.

Next, we assessed whether certain traits shared an evolutionary history with N_2_ fixation gene abundance via phylogenetic linear regression models (**Figure 5, Supplemental Figure 3, Supplemental Tables 5, 11**). Gene abundance (calculated as the proportion of the ninety-three N_2_ fixation machinery genes screened in this study, per genome) was chosen as the independent variable to determine if the complexity of N_2_ fixation genes in a genome could be used to predict other trait values. Response (trait) variables were transformed using either natural log or square root to stabilize variance prior to analysis. Genome size (**Figure 5A, Supplemental Figure 3A**) had a positive, significant linear relationship with N_2_ fixation gene abundance for diazotrophs (*R^2^*=0.104, *P*=2.2×10^-16^). The same trend was observed screening a random sample of non-diazotrophs from our dataset (*R^2^*=0.23, *P*=8.3×10^-42^), suggesting N_2_ fixation gene complexity may interact with broader genomic expansion/subtraction forces, irrespective of an organism’s ability to fix nitrogen.

**Figure 5.**
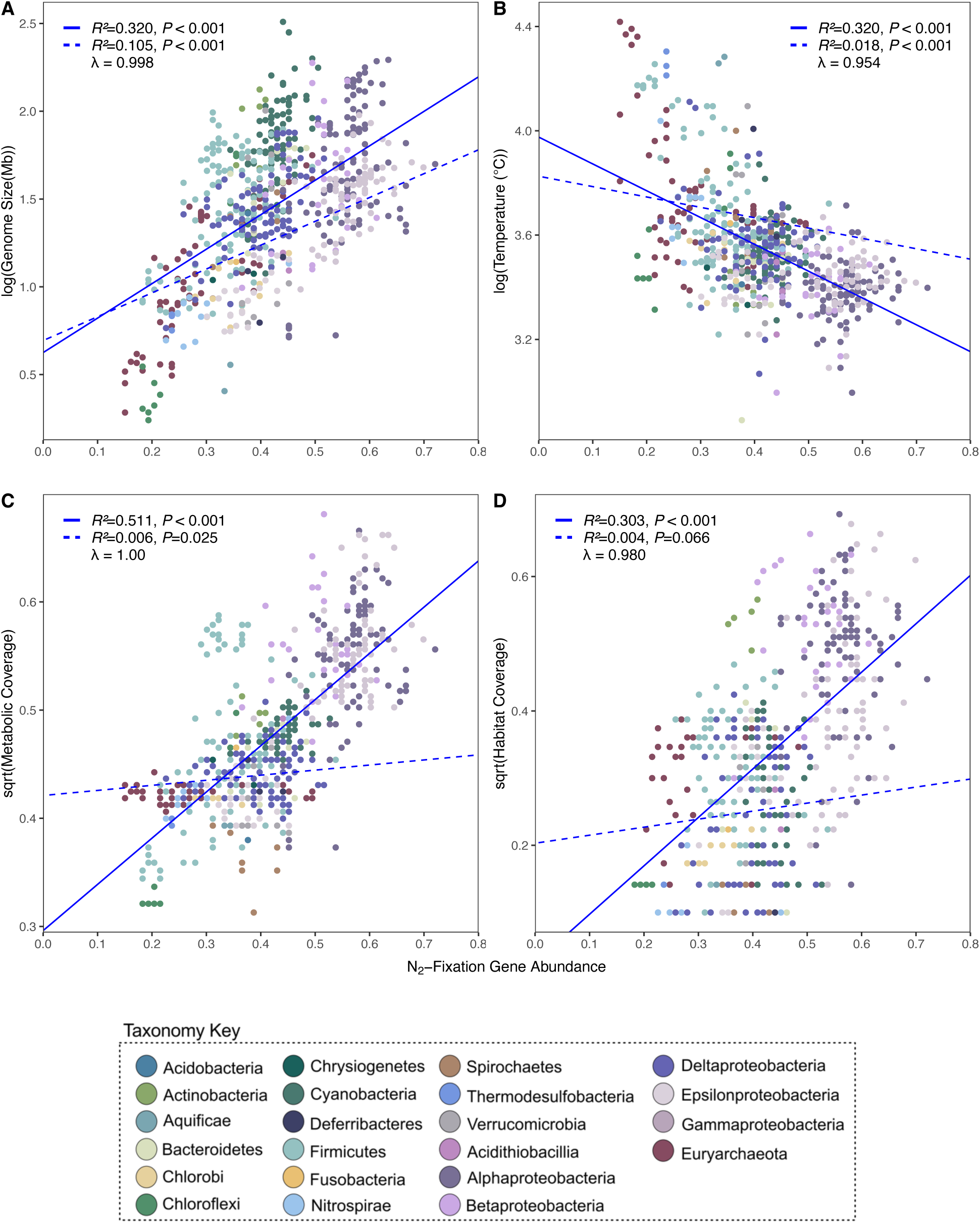
Phylogenetic linear regressions assessed whether the number of N_2_ fixation genes in a genome could predict other traits, such as **A)** genome size, **B)** metabolic coverage, **C)** optimal growth temperature, and **D)** habitat coverage. Positive relationships were observed between N_2_ fixation gene abundance with genome size, metabolic, and habitat coverage. Response variables were either transformed using natural log or square root to stabilize variance prior analysis. Solid blue lines represent non-phylogenetic linear models, dashed blue lines represent phylogenetic linear models. Correlation coefficients (*R^2^*), associated *P*-values for both regression models, and lambda values are shown for each plot. Scatter plots were plotted using ggplot2 in R [56].

For temperature, we initially observed a significant, negative linear relationship with N_2_ fixation gene abundance (**Figure 5B, Supplemental Figure 3B**). Because prior work has shown that adaptation to high temperatures is often linked to genome reduction [55], we repeated the regression using N_2_ fixation gene abundance normalized to genome size (**Supplemental Table 2**). When normalized, the negative relationship with temperature disappeared, suggesting that the original pattern likely reflected genome size constraints rather than a temperature specific effect on N_2_ fixation. A similar pattern emerged for metabolic and habitat coverage. Both traits showed positive relationships with N_2_ fixation gene number, significant for metabolic coverage (*R^2^*=0.006, *P*=0.025) and marginally significant for habitat coverage (*R^2^*=0.004, *P*=0.066) (**Figures 5CD, Supplemental Figures 3CD**). After accounting for genome size, these relationships disappeared, suggesting that gene abundance scales broadly with genomic content rather than specific ecological traits.

### Differences in *N_2_* fixation machinery distribution

We used phylogenetic principal component analysis (PCA) to cluster genomes based on the similarity of their N_2_ fixation gene presence-absence patterns, while accounting for shared evolutionary history. Phylogenetic relationships accounted for most of the variation, as indicated by the low variance captured by the phylogenetic PCA (**Supplemental Figure 10**). This finding suggests that clades are clustered primarily because they share conserved gene sets inherited from common ancestors.

We then examined the key genes driving separation among genomes in the phylogeny-corrected PCA and interpreted their distribution across taxa and oxygen preferences. The ten genes with the strongest influence on each component of the phylogeny-corrected PCA included the different regulation elements (*nifI_2_*, *nifL*, *regRS*, *ntrB*), two different electron-donating/transfer mechanisms (*rnfABCDEG, fixABCX*), nodulation and heterocyst differentiation genes (*nodABCD*, *devC*), oxidative protection mechanisms (*sodN*, *cooA*), cofactor synthesis and iron acquisition genes (*nifM*, *sufD*, *feoC2*), and the genes that make up both the Fe- and V-nitrogenases (*vnfHGDKE*, *anfHGDK*). The post- translational *nifI_2_* regulator is primarily found in anaerobes, such as Archaea and Firmicutes [12, 13], while other regulatory mechanisms such as *regS*, *nifL*, and *ntrB* are primarily found in aerobic/facultative Proteobacteria (**Supplemental Figure 1**). Pearson correlations of gene co-occurrence (**Supplemental Figure 11**) confirmed that *nifI_2_* and *nifI_1_* had strong negative correlations with genes *nifTQWXZ*, *fnr*, *fdx*, *oxyR,* and *ntrBC*, many of which were correlated with the ability to use oxygen in diazotrophs [12].

A preference for either *rnfACDEG* or *fixACX* electron donation systems was evident in the PCA where the genes pointed opposite from one another (**Supplemental Figure 10,11**). Nodulation genes *nodABC* also pointed opposite to *rnfACDEG* and instead had a positive correlation with *fixACX* (**Supplemental Figure 11**). This is confirmed by the fact that diazotrophs forming symbioses with plants through nodulation prefer *fix* genes over *rnf* genes for electron donation [57, 58]. None of the N_2_ fixation genes were unique to thermophiles and neither Fe- nor V- nitrogenases were found in genomes with optimal growth temperatures greater than 41°C.

### Dynamic history of gene gain and loss

We estimated net gain or loss of N_2_ fixation machinery genes at each node of the nitrogenase-containing species tree using Wagner parsimony (see methods). Our results show a dynamic history of gene gain and loss over time (**Figure 6**). Specifically, net gene gain was predicted at many of the early branching nodes, whereas gene loss primarily occurred closer to terminal tree branches. This suggests an initial expansion of genes early in diazotroph evolution, followed by successive gene loss events, potentially corresponding to an expansion of diazotroph lineages into new environments where certain components of the N_2_ fixation machinery became obsolete.

**Figure 6.**
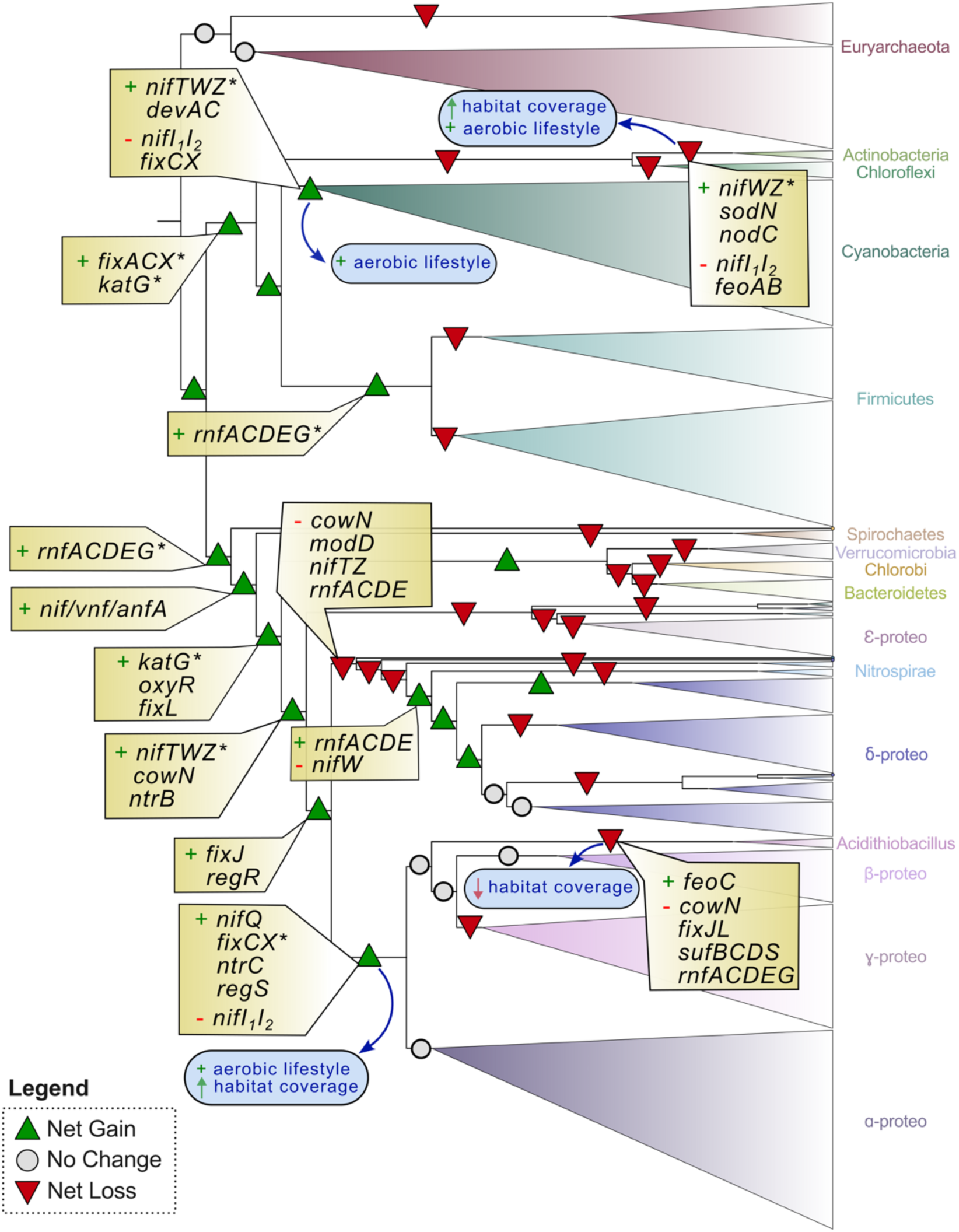
N_2_ fixation gene gain (green triangles) and gene loss (red triangles) throughout the evolution of diazotrophs. Events are mapped to the WoL [20] species tree which was pruned to include only tips associated with diazotrophs identified in this study (n=673). Gene gain dominated the earliest nodes, whereas gene loss was most common near terminal branches. No change (grey circle) represents no gain or loss at that node, or an equal number or gains and losses. Asterisks highlight potential candidates for HGT or genes derived from non-diazotroph ancestors not in this subsetted phylogeny. All gene events can be found in Supplemental table 8. Key transitional events of gene abundance and habitat coverage can be found in Supplemental Figure 12.

The nodes with the highest number of gene gains (+11) occurs at the most recent common ancestor (MRCA) of Bacteria and at the MRCA of the Euryarchaeota clade Methanosarcina (not shown). Some of the clades that experienced large gene losses include Acidithiobacillia (-18), the Firmicute-Clostridia clade Thermoanaerobacterales (- 18) (not shown), Thermodesulfobacteria (-17) (not shown), Nitrospirae (-16), and the MRCA of Aquificae and Epsilonproteobacteria (-11) (**Supplemental Tables 7-8**). Acidothiobacillia’s loss of genes *cowN*, *fixJL*, and the Suf and Rnf systems occurs in tandem with a loss of habitat coverage (**Figure 6**). The loss of post-transcriptional regulators *nifI_1_I_2_* unsurprisingly coincides with a gain in aerobic lifestyles at the MRCA of Cyanobacteria, Actinobacteria and the MRCA of Gamma- and Alphaproteobacteria (**Figure 6**) as *nifI_1_I_2_* is the main regulatory mechanism in anaerobes. The appearance of transcriptional regulation by *nifA*, mostly used by aerobes, predates the loss of *nifI_1_I_2_* in the MRCA of Gamma- and Alphaproteobacteria (**Figure 6**). The gain of genes involved in protection against oxidants (*katG*, *oxyR*, and *cowN*) and oxygen response regulators *fixJL* occurs early along the MRCA nodes of Verrucomicrobia, Chlorobi, Bacteroidetes, and Proteobacteria (**Figure 6**). We noted two distinct gains of *rnfACDEG*, *nifTWZ*, *katG* and *fixCX* in distantly related diazotrophs, indicating they were either inherited from a common ancestor that does not have nitrogenase (and therefore would not be represented in this tree), and/or they were gained through horizontal gene transfer.

## Discussion

### Nitrogenase Distribution, Diversity, and Evolution Across Prokaryotic Lineages

Nitrogenase is found across a taxonomically diverse range of microbes with ecologies spanning root-associated symbiosis in the rhizosphere [59] to deep-sea sediments kilometers below the ocean surface [60–62]. The factors underlying this broad yet uneven distribution remain unclear but likely reflect a complex evolutionary history shaped by dynamic interplay between nitrogenase, its host cell, and the surrounding environment.

To investigate these patterns, we developed a comprehensive catalog of 673 nitrogenase-encoding genomes and their associated N_2_ fixation machinery, tracing historical genomic changes that shaped the evolution of this important metabolism (**Figure 3**). In addition to the *nif* cluster and its accessory genes, we included 63 genes outside of this cluster with reported roles in N_2_ fixation, yielding a total of 93 genes representing the N_2_ fixation genetic machinery (**Table 1, Supplemental Table 1**). These data were integrated with host-associated trait information (e.g., oxygen metabolism, temperature, pH, genome size, habitat coverage), linking the evolutionary history of nitrogenase and its supporting genes to the environmental diversity of N_2_ fixation observed today.

Oxygen inhibits nitrogenase activity [63], and modern diazotrophs employ a range of strategies to protect the enzyme from this interference [19, 64, 65]. Consistent with previous findings [12], we observed that on average, aerobic diazotrophs harbor more N_2_ fixation-associated genes than anaerobes (**Figure 4B**), likely reflecting the additional genetic components required for oxygen protection and regulation under aerobic conditions. For example, we found that genes *katG*, *oxyR*, and *cowN*, which provide protection against oxidants [66, 67], along with the oxygen response regulators *fixJL* [15] were acquired along the lineages leading up to the divergence of Proteobacteria (**Figure 6**), a period that coincides with the historic accumulation of oxygen in the atmosphere ∼2.45 billion years ago known as the Great Oxidation Event (GOE) [68]. This pattern suggests these genes played a role enabling ancient diazotrophs to expand into increasingly oxygenated environments. Recent studies further support the divergence of the alternative nitrogenases during this transition to an oxic atmosphere [69], underscoring how environmental transitions, particularly the historical rise of atmospheric oxygen levels, reshaped the molecular evolution of N_2_ fixation in deep time.

We recapitulate the relationship between N₂ fixation gene abundance and temperature previously reported by Boyd et al. [12], that on average, thermophiles possess the simplest N_2_ fixation-related gene sets (**Figure 4AB**, **Figure 5B**). Our analyses indicate that this pattern is in part due to genome reduction associated with long-term adaptation to high temperatures [55]. Yet, if the earliest diazotrophs were indeed thermophilic [12], then temperature’s role shaping the evolution of N_2_ fixation cannot be dismissed. For example, some cold-adapted diazotrophs switch from Mo- to V-based nitrogenases [70–72], indicating how shifts in temperature regimes may have shaped the functional diversification of nitrogenase over evolutionary time.

Outside our set of nitrogenase-containing genomes, we identified genomes with partial detection of the core N_2_ fixation genes. Notably, several anaerobic thermophiles lack *nifE*, *nifN*, or both (**Figure 3**). These genes are involved in FeMo cofactor assembly and are part of a proposed minimal *nif* gene set (*nifHDKENB)* required for N_2_ fixation [73]. Lineages exhibiting this partial gene set include Euryarcheaota (such as members of the ANME groups [74]), Chloroflexi, and Firmicutes. Some have verified N_2_ fixation activity, while others, like *Roseiflexus castenholzii*, do not [75–78]. Ancestral sequence reconstructions suggest that *nifDK* may have originated from a duplicated maturase-like *nifEN* ancestor [79], raising the possibility that some extant diazotrophs lacking *nifEN* evolved alternative cofactor maturation strategies, perhaps as a way to maximize energy conservation under thermophilic conditions [55]. Support for this idea comes from *Roseiflexus* sp. RS-1, which lacks *nifEN* and appears to use *nifDK* to fulfill both the cofactor maturase and catalytic roles [80]. Further work will be required to determine whether this streamlined mechanism occurs in other diazotrophs missing *nifEN*.

Plant-associated genomes showed the highest N_2_ fixation gene counts (**Figure 4C**), consistent with the pivotal role of rhizobia in plant host nitrogen supply [81, 82]. Our reconstructions capture the divergence of Actinobacteria *Frankia* from its ancestor, marked by a transition to aerobic metabolism, accompanying genome and habitat expansion, the loss of *nifI_1_I_2_* regulators, and the gain of both nodulation (*nodC*) and oxidative stress response (*sodN*) genes to support its dual free-living and symbiosis states (**Figure 6**).

We observed that *Frankia* lacks key regulators such as *nifAL* [17]. However, we noted the presence of the post-translational regulator *draG*, without its partner *draT*, indicating that draG-draT regulation is likely non-functional and perhaps a novel, as-yet-unknown mechanism for nitrogenase inactivation is present [83] – a possibility that warrants further study.

In many plant symbioses, oxygen levels are tightly regulated by the host to activate the FixL - FixJ pathway, ultimately driving nitrogenase expression [84], while *nod* genes mediate nodulation [85]. Terrestrial habitats, being more variable than oceans [86], impose fluctuating environmental conditions that demand dynamic nitrogenase regulation [87–89]. Soil molybdenum scarcity likely favors the deployment of alternative nitrogenases [90], contributing to the expansion of gene repertoire in terrestrial environments. Our results illustrate this pattern. For example, several lichen-symbionts within the Nostocales order of Cyanobacteria possess V-nitrogenase [91], likely as a way to cope with limited Mo in high latitude boreal forests [92]. Together, these findings highlight how diazotroph N_2_ fixation gene repertoires have been shaped by environmental constraints and the selective pressures of symbiotic lifestyles.

### Eco-evolutionary history of nitrogenase-associated genes

Central to our framework is that N_2_ fixation gene complexity is not an isolated trait, rather, it is embedded within a larger evolutionary framework of genome expansion, metabolic versatility, and habitat diversity (**Figures 4, 5**). Gene complexity can even serve as an ecological marker distinguishing generalists, with broad habitats and large, versatile genomes, from specialists with narrow niches and smaller genomes [93–95]. This distinction is particularly relevant today as specialist diazotrophs may contribute disproportionately to total soil N_2_ fixation and thereby differentially influence nitrogen input into these environments [96].

Our reconstructions show that as diazotrophs diversified, so did their N_2_ fixation machinery (**Figure 3**). Gene gain dominated early evolution, with clades such as Alpha-, Gamma-, and Betaproteobacteria maintaining extensive N_2_ fixation genes (**Figure 6**). In contrast, later stages were marked by lineage-specific losses, particularly in terminal branches, suggesting a shift towards ecological specialization and genome streamlining in diazotrophs. Losses were most pronounced in Chloroflexi, Nitrospirae, Aquificae, Acidithiobacillia, Thermodesulfobacterota, and Epsilonproteobacteria, groups often associated with anaerobic, thermophilic, acidophilic, or host-adapted lifestyles (**Supplemental Table 2**).

In some cases, N_2_ fixation gene loss tracked closely with reduced habitat coverage, as in Epsilonproteobacteria and Acidithiobacillia (**Figure 6**), consistent with prior work on genome streamlining in these clades [97–99]. Adaptation to deep-sea hydrothermal vents [100], shifts toward heterotrophy [97], and transitions to acidophily [101] all show how niche transitions often coincide with simplification of N_2_ fixation gene sets. There are exceptions to this trend between genome size and the size of the N_2_ fixation machinery. For example, *Zymomonas mobilis* (Alphaproteobacteria) maintains the highest proportion of N_2_ fixation genes (∼2-3% of all coding genes) despite having a compact ∼2Mb genome (**Supplemental Table 2**). Similarly, in symbiont systems, genome reduction often reflects host compensation [102]. UCYN-A2, a marine symbiont recently identified as an early stage “nitroplast” in *Braarudosphaera bigelowii* [103, 104], has lost CO_2_ fixation and photosystem II [105, 106], yet retains an intact *nif* cluster (*nifTZVBSHDKENXW)* [107].

Horizontal gene transfer (HGT) has further shaped N_2_ fixation evolution [11, 108], likely explaining several gene acquisitions we observed (e.g., *rnfACDEG*, *nifTWZ*, *katG* and *fixCX* in **Figure 6**). Previous studies have documented entire *nif* clusters moving across major clades – including a >20kb transfer from Gammaproteobacterium in to the diatom symbiont “*Candidatus* Tectiglobus diatomicola” [86, 109]. In our dataset, *Coleofasciculus chthonoplastes* PCC 7420 (formerly *Microcoleus chthonoplastes*) appears to have acquired *nifHDKEN* from a Deltaproteobacterium [88, 110]. Together, these patterns show that the complexity of N_2_ fixation genes reflects a balance between the forces of gene loss and gain, specialization, horizontal transfer - all embedded in the broader trajectory of diazotroph genome evolution.

### Limitations of comparative genomics

As with any comparative genomic approach, there are inherent limitations in identifying the full complement of genes and traits within genomes. One potential limitation is that our analysis focuses on genes already recognized to be playing a role in N_2_ fixation, meaning additional, yet-undiscovered genes may contribute to the process, particularly in non-model diazotrophs, where many functional mechanisms remain uncharacterized. Another caveat is our reliance on KEGG KO number. Not all genes could be assigned KO numbers, and some KO identifiers fail to distinguish between genes with high sequence similarity, such as *nifA*/*vnfA*/*anfA*. Continued improvements in gene functional annotation and database curation will be essential for building a more complete understanding of diazotroph capabilities [111].

## Conclusion

Examining the distribution of diverse N_2_ fixation machinery offers insight into which genes were critical for enabling transitions into new environments throughout evolutionary history. The rise of atmospheric oxygen, for example, fundamentally reshaped electron-donor availability [112] and, together with changes in metal availability [113, 114], likely exerted strong selective pressures on N_2_ fixation systems, as demonstrated here.

Reconstructing historical patterns from modern genetic records is thus pivotal for understanding the processes that govern evolution [115]. Global trends in genome evolution not only reveal how organisms responded to past environmental fluctuations but connects them to their present-day ecological niches.

In his book *Life’s Engines*, Paul Falkowski likens nitrogenase and its core machinery to a computer’s hardware, while accessory genes function as the software, essential for maintenance, optimization and updates [116]. This analogy is compelling, capturing the deep interconnection between core biochemical processes and their regulatory networks. These elements must evolve in concert to navigate the challenges posed by changing environments, a dynamic reflected in the expansions and contractions within diazotroph genomes which we have documented across both ancient and modern contexts.

## Acknowledgements

We gratefully acknowledge support from the NASA Interdisciplinary Consortium for Astrobiology Research (ICAR): Metal Utilization and Selection Across Eons, MUSE under grant no 80NSSC17K0296, and the NASA Exobiology Program under grant no NNH23ZDA001N. M.S. was supported by the NASA Postdoctoral Program, administered by Oak Ridge Associated Universities under contract with NASA. We thank Patricia Tran, Shaomei He, colleagues in the Kaçar Lab, members of NASA MUSE ICAR, and colleagues at Texas State University for helpful feedback on manuscript; and the Center for High Throughput Computing (CHTC) and the Department of Bacteriology at the University of Wisconsin–Madison for providing computing resources. We would like to finally thank the four anonymous reviewers for their constructive feedback.

## Data availability

All custom scripts, tree files, raw data, and PDF versions of the figures are available at https://github.com/kacarlab/Sobol_2025_NifGeneEcoEvo. All supplemental tables are available in the uploaded datasets. Specifically, supplementary Tables 2 and 9 include genome accession numbers, taxonomy, and trait data information.

## Notes

https://github.com/kacarlab/Sobol_2025_NifGeneEcoEvo.

### Competing Interest Statement

The authors have declared no competing interest.

### Summary of Updates

New analyses and accompanying figure, updated SI, updated text for clarity.

