## Supplemental Information for "Ecological constraints and evolutionary trade-offs shape nitrogen fixation across habitats"

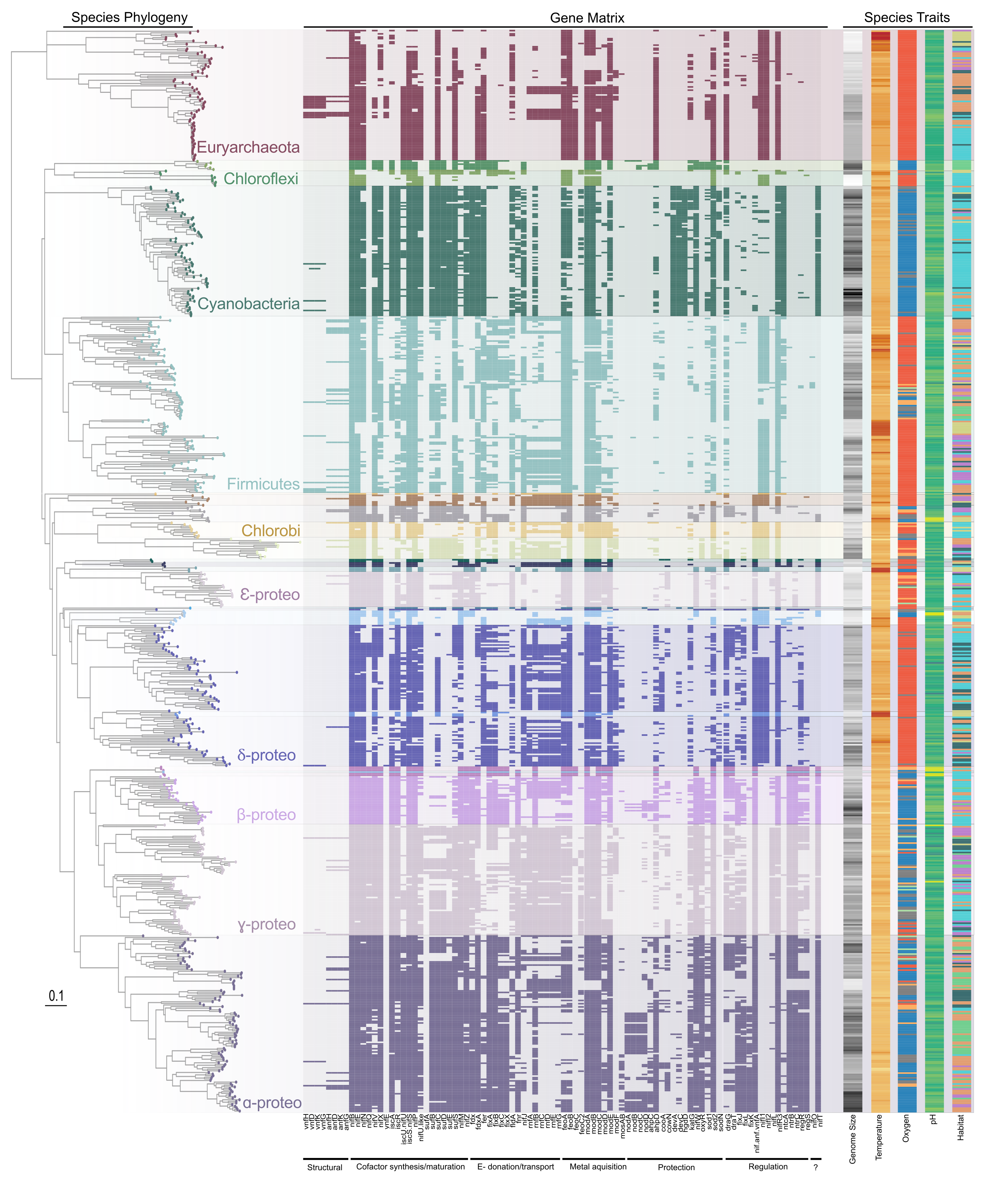

**Supplemental Figure 1.** Presence-absence distribution N_2_ fixation genes amongst genomes that have nitrogenase in the Zhu et al (2020) Web of Life (WoL) phylogeny [1]. This figure accompanies Figure 1 in the main manuscript. Genes are organized into categories based on their functional role (structural, cofactor synthesis, electron donation/transport, metal acquisition, protection, regulation, and unknown). The “?” indicates genes *nifO* and *nifT* whose functions are unknown. The gene presence-absence matrix was determined via homology search with HMMER against the KEGG KO database [2]. Species-associated ecological and physiological traits were mapped alongside the phylogeny and matrix to highlight associations in genes and gene abundance with different lifestyles. The list of genes and their roles can be found in Supplemental Table 1, and the gene matrix and associated categories can be found in Supplemental Table 3. The phylogeny and matrices were plotted using the ggTree R package [3].

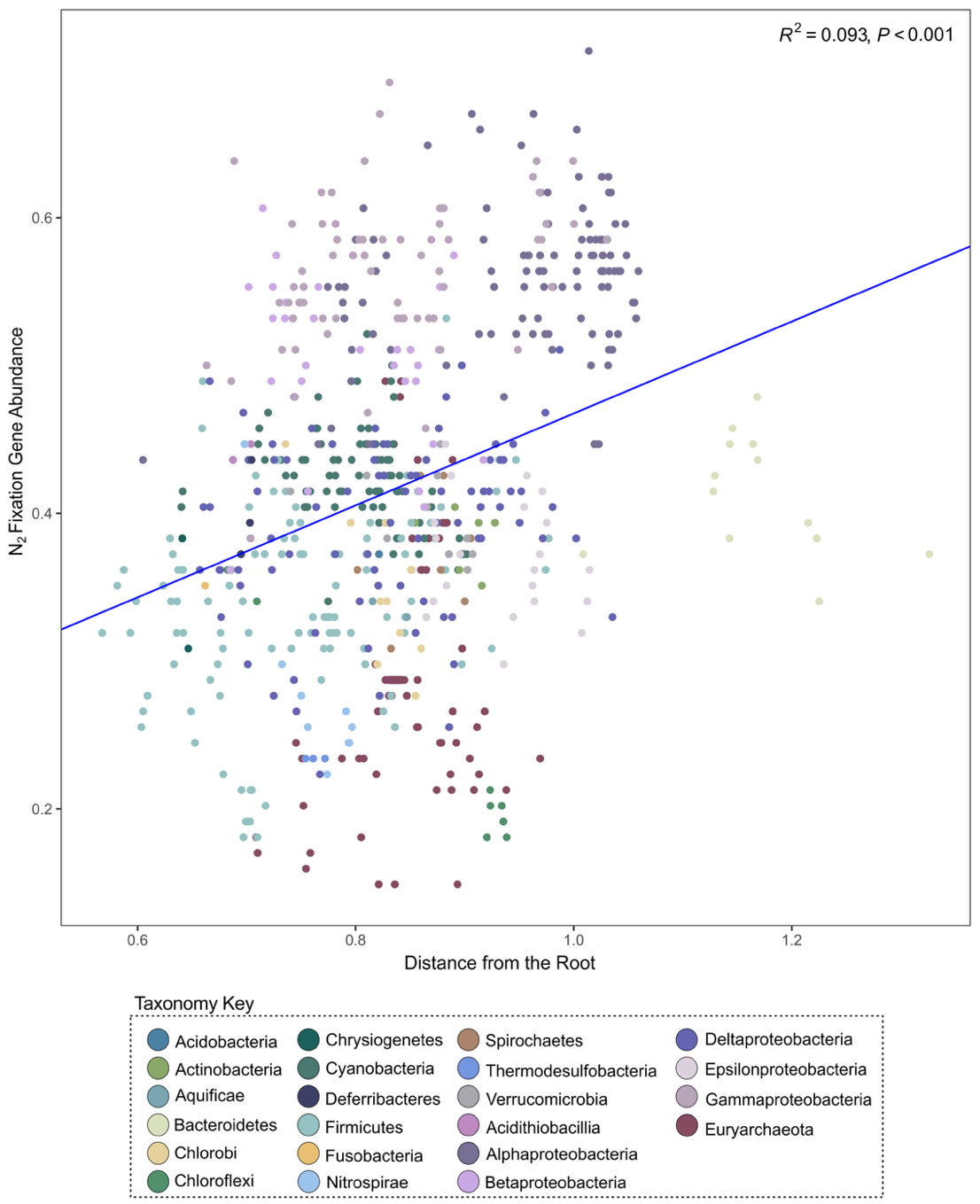

**Supplemental Figure 2.** Relationship between patristic distances and N_2_ fixation gene abundance. A positive relationship was observed, indicating that later diverged taxa have significantly (p<0.001) larger N_2_ fixation gene repertoires. Each point represents a genome, colored by taxonomic affiliation. The linear regression and trendline were calculated using the *lm* function in the stats R package [4]. The *distRoot* function in the **adephylo** R package [5] was used to calculate the root-to-tip patristic distances. The scatter plot was plotted using ggplot2 in R [6].

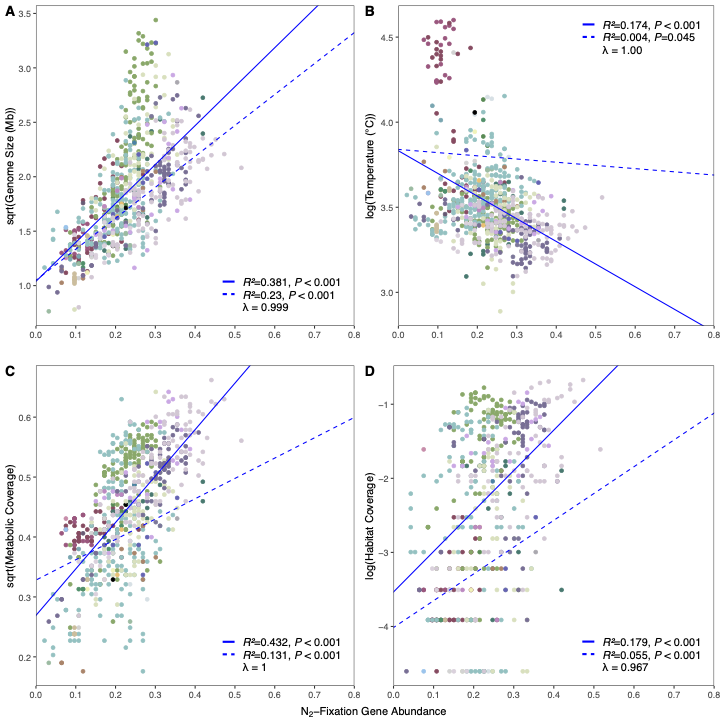

**Supplemental Figure 3.** Phylogenetic linear regressions assessing whether the abundance of N_2_ fixation-related genes can predict different traits of non-diazotroph hosts. The different traits assessed are shown on the y-axes and include genome size (A), temperature (B), metabolic coverage (C), and habitat coverage (D). Response variables were either transformed using natural log or square root to stabilize variance prior analysis. Solid blue lines represent non-phylogenetic linear models, dashed blue lines represent phylogenetic linear models. Correlation coefficients (*R^2^*) and associated *P*-values for both regression models, and lambda values are shown for each plot. Scatter plots were plotted using ggplot2 in R [6]. Points are colored by taxonomy following the color scheme in the legend of Figure 5 in the main text.

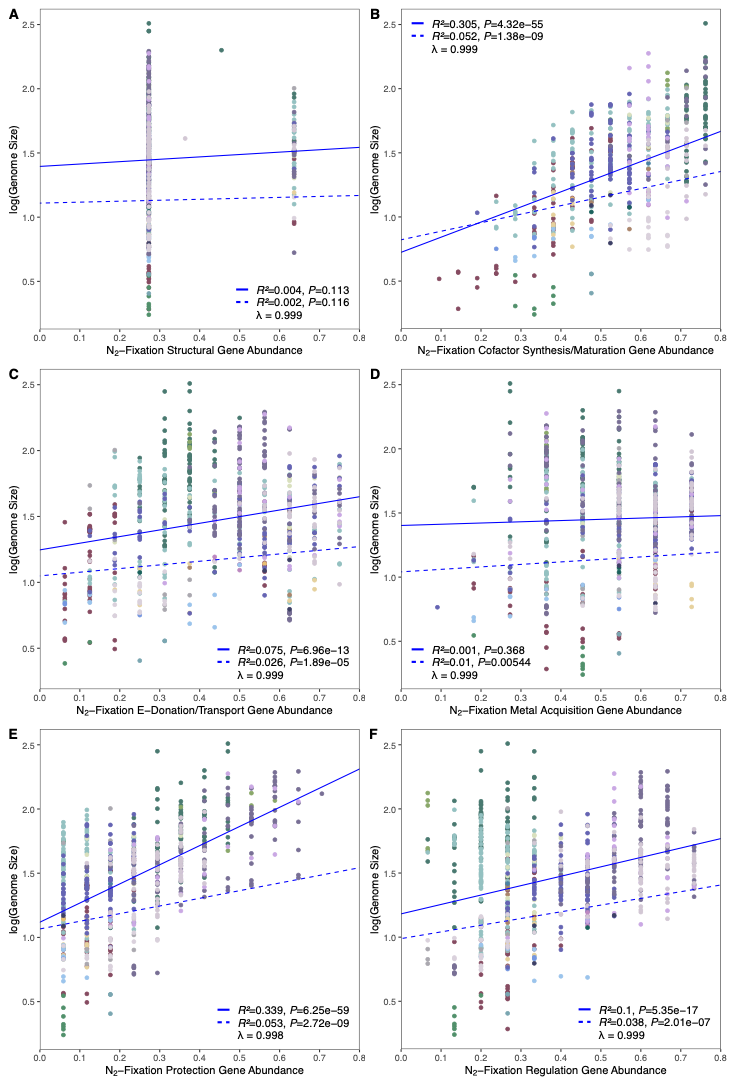

**Supplementary Figure 4.** Phylogenetic linear regressions assessing whether the abundance of N_2_-fixation genes belonging to different functional groups can predict the size of genomes of diazotrophs. Functional groups include those related to nitrogenase structure (A), cofactor synthesis/maturation (B), electron donation/transport (C), metal acquisition (D), protection (E), and regulation (F). Response variables were either transformed using natural log or square root to stabilize variance prior analysis. Solid blue lines represent non-phylogenetic linear models, dashed blue lines represent phylogenetic linear models. Correlation coefficients (*R^2^*) and associated *P*-values for both regression models, and lambda values are shown for each plot. Scatter plots were plotted using ggplot2 in R [6]. Points are colored by taxonomy following the color scheme in the legend of Figure 5 in the main text.

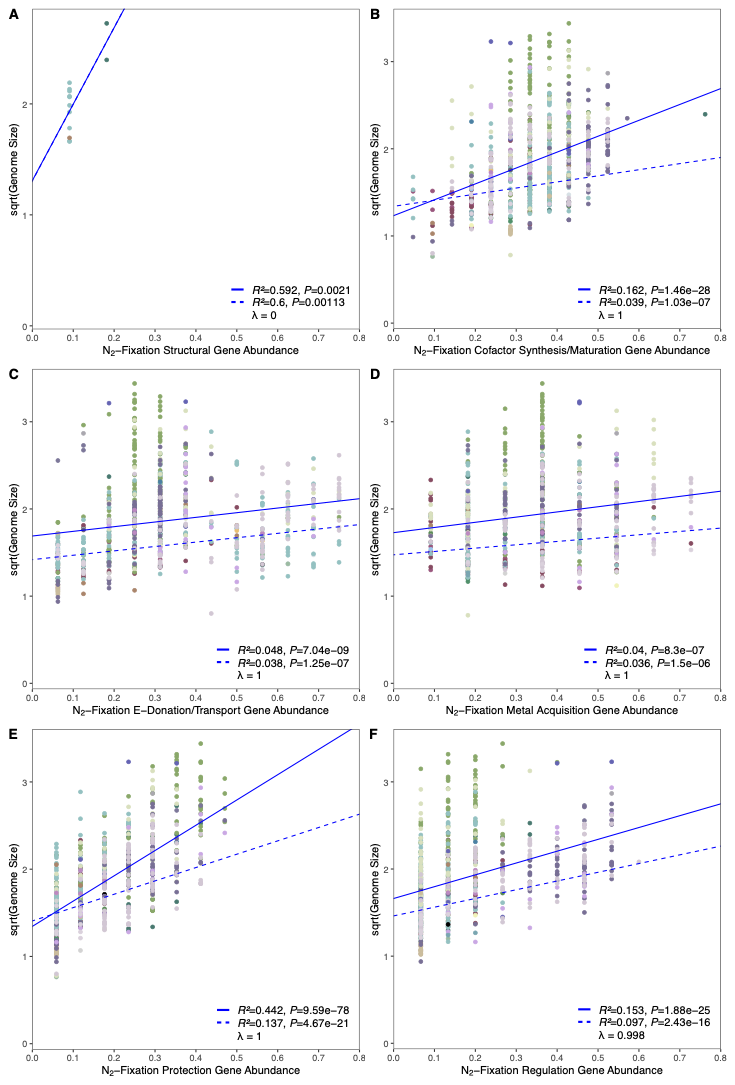

**Supplementary Figure 5.** Phylogenetic linear regressions assessing whether the abundance of N_2_-fixation genes belonging to different functional groups can predict the size of genomes of non-diazotrophs. Functional groups include those related to nitrogenase structure (A), cofactor synthesis/maturation (B), electron donation/transport (C), metal acquisition (D), protection (E), and regulation (F). Response variables were either transformed using natural log or square root to stabilize variance prior analysis. Solid blue lines represent non-phylogenetic linear models, dashed blue lines represent phylogenetic linear models. Correlation coefficients (*R^2^*) and associated *P*-values for both regression models, and lambda values are shown for each plot. Scatter plots were plotted using ggplot2 in R [6]. Points are colored by taxonomy following the color scheme in the legend of Figure 5 in the main text. All correlations for phylogenetic regressions are notably weak, suggesting phylogenetic relationships may be an important factor driving the observed trends between traits and gene abundance. One exception is the correlation between genome size and protection gene abundance which shared a positive correlation (*R^2^*=0.137, *P*=4.67×10^-21^), suggesting oxygen protection genes may have been more involved in genome expansions/subtractions in non-diazotrophs, compared to genes from other functional groups.

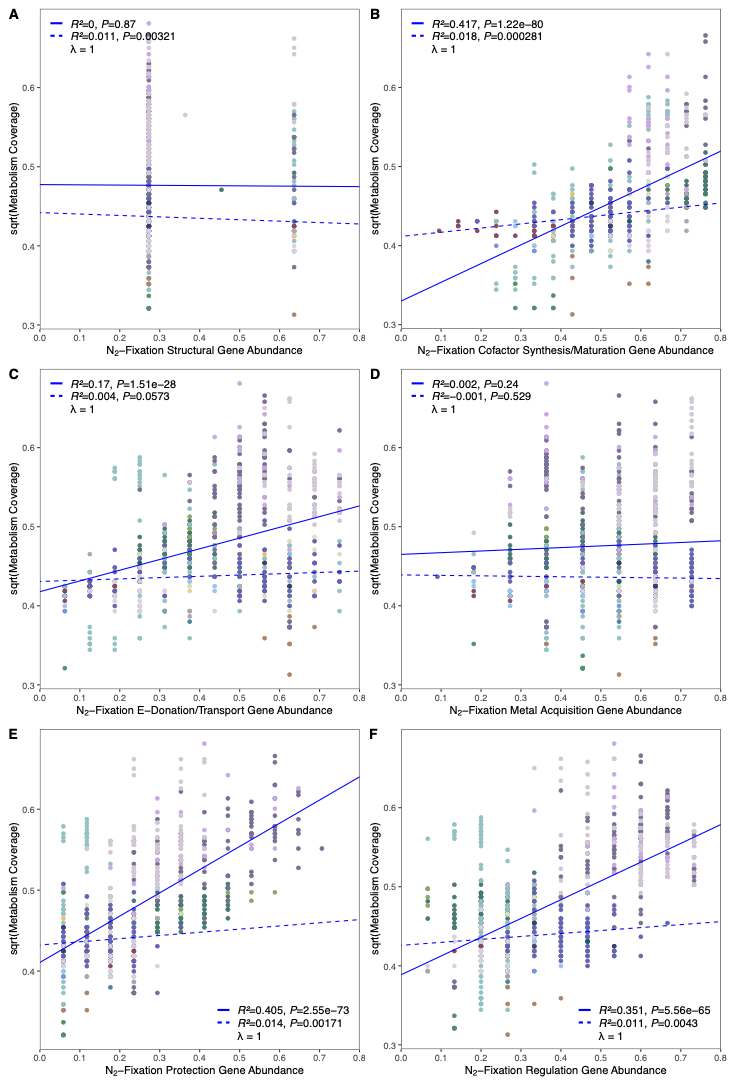

**Supplementary Figure 6.** Phylogenetic linear regressions assessing whether the abundance of N_2_-fixation genes belonging to different functional groups can predict metabolic coverage of diazotrophs. Functional groups include those related to nitrogenase structure (A), cofactor synthesis/maturation (B), electron donation/transport (C), metal acquisition (D), protection (E), and regulation (F). Response variables were either transformed using natural log or square root to stabilize variance prior analysis. Solid blue lines represent non-phylogenetic linear models, dashed blue lines represent phylogenetic linear models. Correlation coefficients (*R^2^*) and associated *P*-values for both regression models, and lambda values are shown for each plot. Scatter plots were plotted using ggplot2 in R [6]. Points are colored by taxonomy following the color scheme in the legend of Figure 5 in the main text.

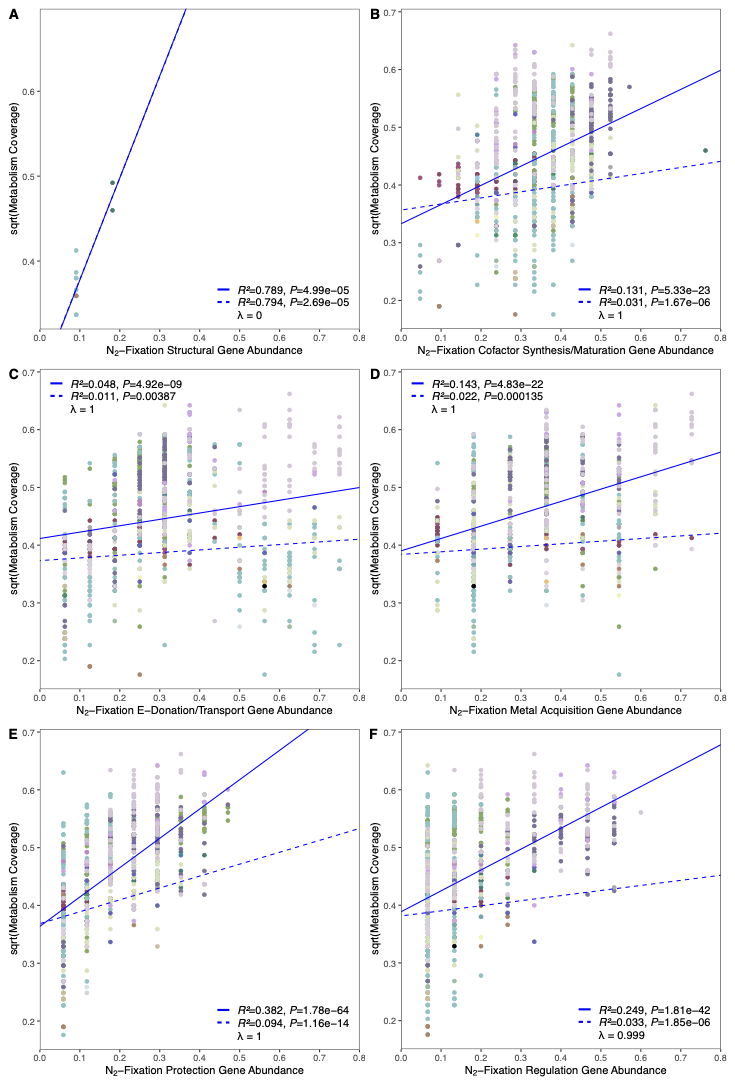

**Supplementary Figure 7.** Phylogenetic linear regressions assessing whether the abundance of N_2_-fixation genes belonging to different functional groups can predict metabolic coverage of non-diazotrophs. Functional groups include those related to nitrogenase structure (A), cofactor synthesis/maturation (B), electron donation/transport (C), metal acquisition (D), protection (E), and regulation (F). Response variables were either transformed using natural log or square root to stabilize variance prior analysis. Solid blue lines represent non-phylogenetic linear models, dashed blue lines represent phylogenetic linear models. Correlation coefficients (*R^2^*) and associated *P*-values for both regression models, and lambda values are shown for each plot. Scatter plots were plotted using ggplot2 in R [6]. Points are colored by taxonomy following the color scheme in the legend of Figure 5 in the main text.

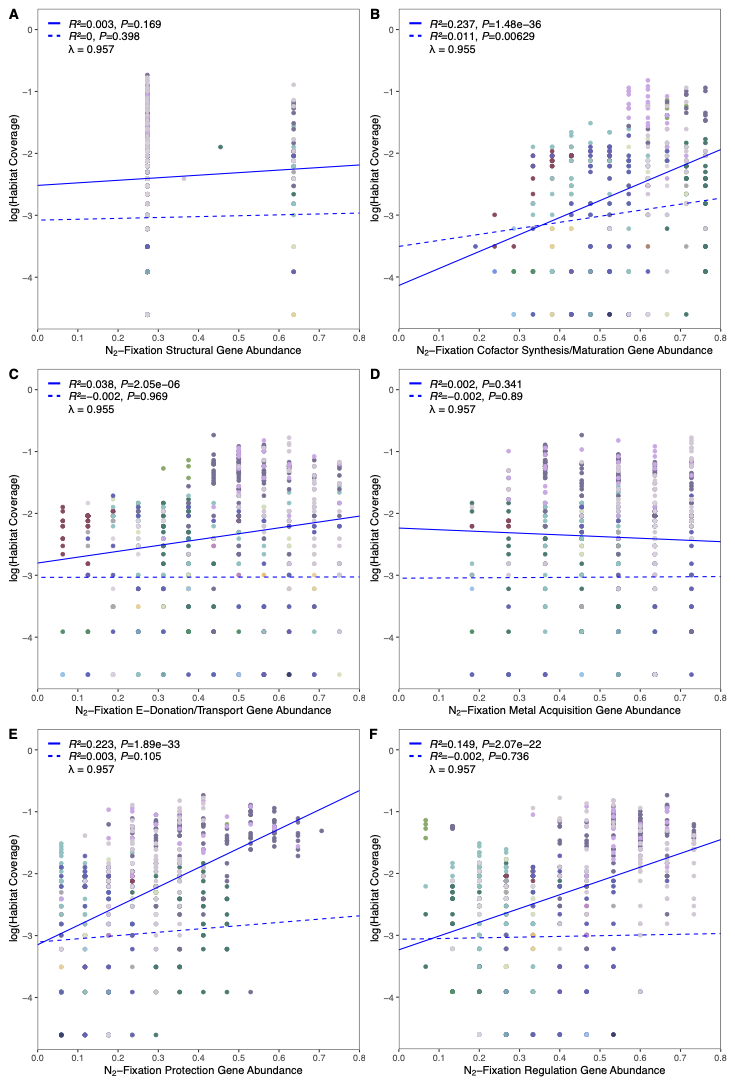

**Supplementary Figure 8.** Phylogenetic linear regressions assessing whether the abundance of N_2_-fixation genes belonging to different functional groups can predict habitat coverage of diazotrophs. Functional groups include those related to nitrogenase structure (A), cofactor synthesis/maturation (B), electron donation/transport (C), metal acquisition (D), protection (E), and regulation (F). Response variables were either transformed using natural log or square root to stabilize variance prior analysis. Solid blue lines represent non-phylogenetic linear models, dashed blue lines represent phylogenetic linear models. Correlation coefficients (*R^2^*) and associated *P*-values for both regression models, and lambda values are shown for each plot. Scatter plots were plotted using ggplot2 in R [6]. Points are colored by taxonomy following the color scheme in the legend of Figure 5 in the main text.

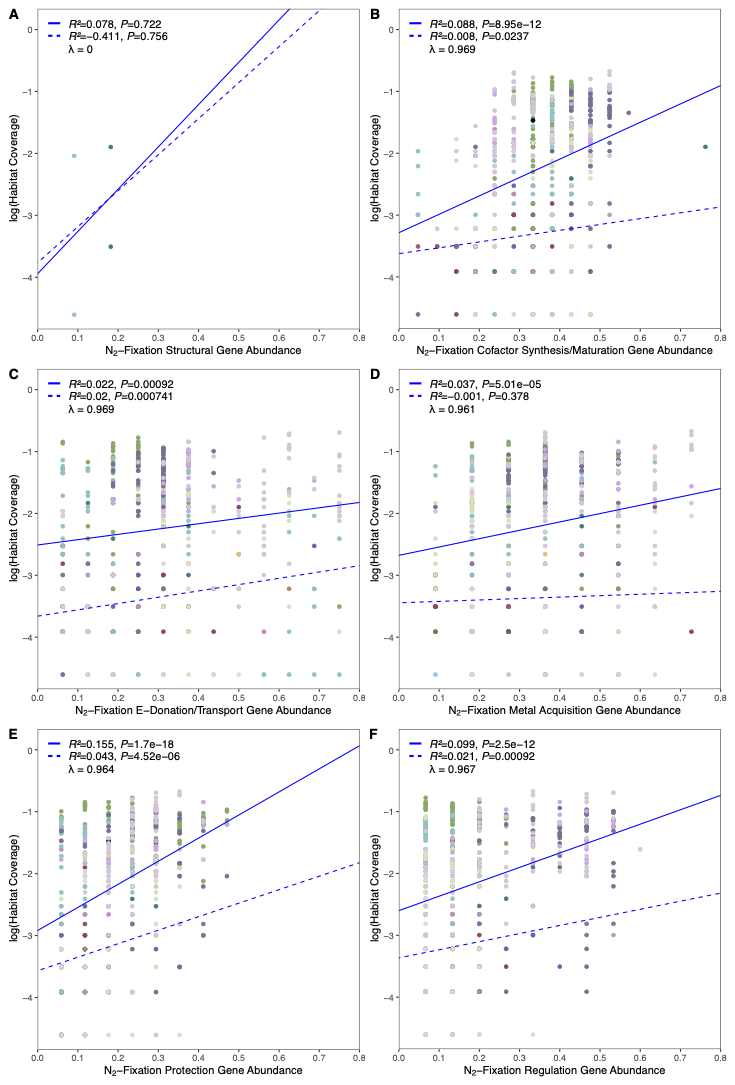

**Supplementary Figure 9.** Phylogenetic linear regressions assessing whether the abundance of N_2_-fixation genes belonging to different functional groups can predict habitat coverage of non-diazotrophs. Functional groups include those related to nitrogenase structure (A), cofactor synthesis/maturation (B), electron donation/transport (C), metal acquisition (D), protection (E), and regulation (F). Response variables were either transformed using natural log or square root to stabilize variance prior analysis. Solid blue lines represent non-phylogenetic linear models, dashed blue lines represent phylogenetic linear models. Correlation coefficients (*R^2^*) and associated *P*-values for both regression models, and lambda values are shown for each plot. Scatter plots were plotted using ggplot2 in R [6]. Points are colored by taxonomy following the color scheme in the legend of Figure 5 in the main text.

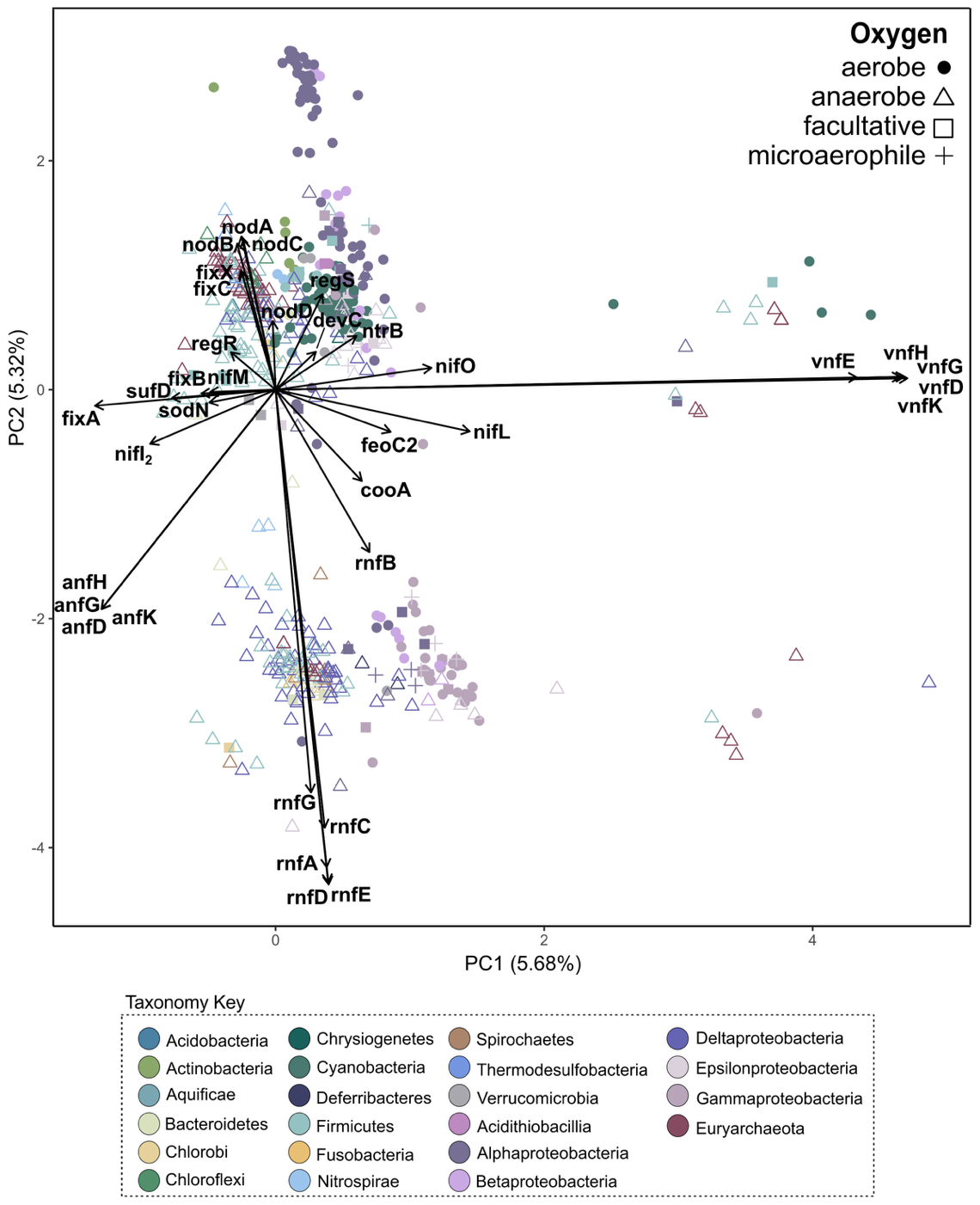

**Supplemental Figure 10.** Phylogenetic principal component analysis (PCA) of genomes clustered by their N_2_ fixation gene presence-absence patterns (Supplemental Figure 1), performed with the *phyl.pca* function (method = “BM”, mode =”corr”) in phytools R package [7]. Each point represents a genome, colored by taxonomic affiliation, and shapes represent oxygen preference. Vectors indicate the top ten genes with the strongest influence on each component. The low explained variance of the first two components (PC1 = 5.68%, PC2 = 5.32%) suggests that most variation is explained by phylogenetic structure such that genomes cluster primarily due to conserved gene sets inherited from common ancestors. The PCA was plotted using ggplot2 in R [6].

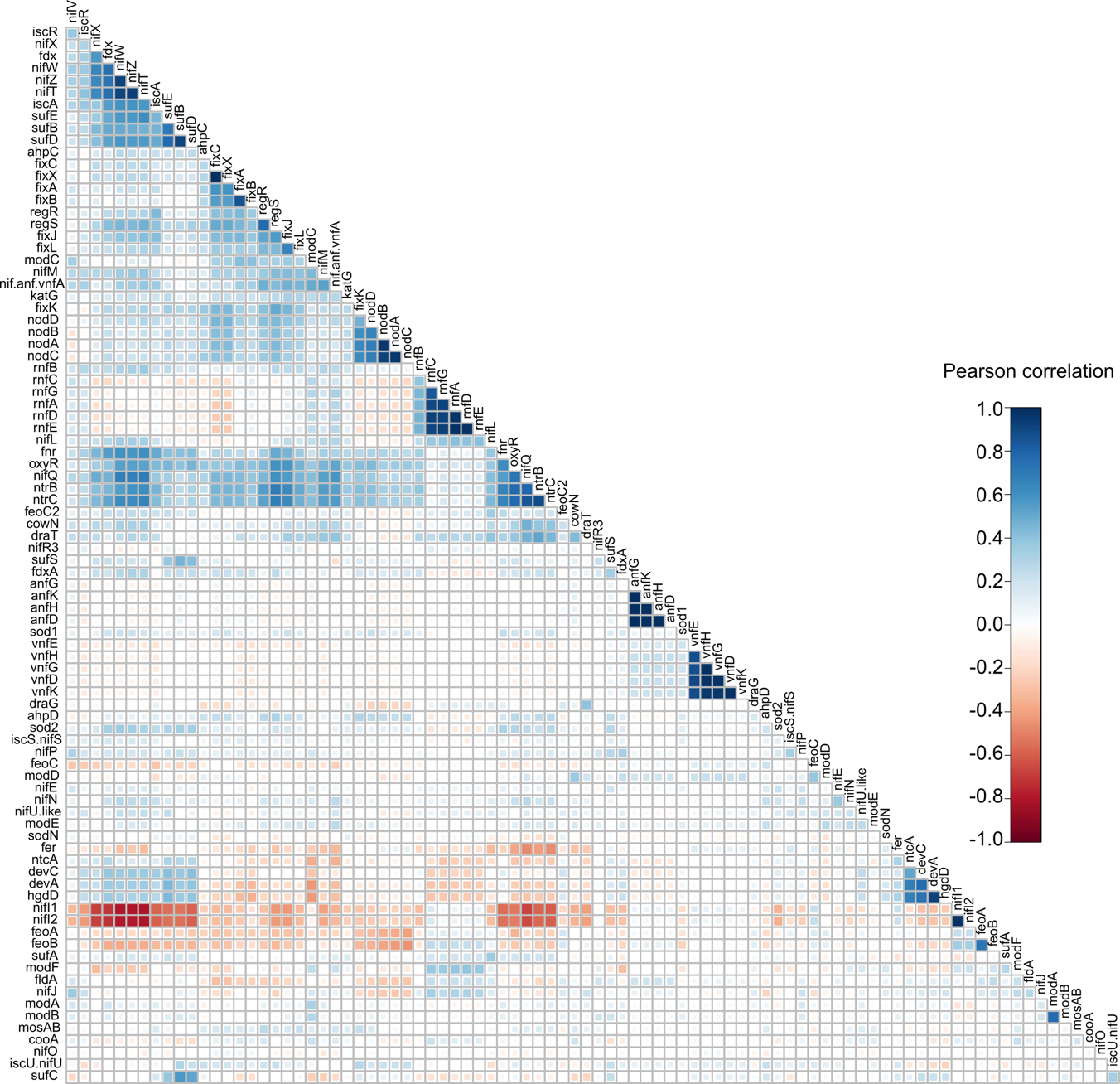

**Supplemental Figure 11.** Pearson correlation of gene co-occurrence among N_2_ fixation genes, based on the presence-absence matrix in Supplemental Figure 1. Each cell represents the correlation between a pair of genes, with blue indicating positive correlation (+1) and red indicating a negative correlation (-1). Correlations were calculated with the *cor* function in the stats R package [4] and plotted with corrplot [8] using hierarchical clustering. Non-significant correlations (p>0.05) were left blank.

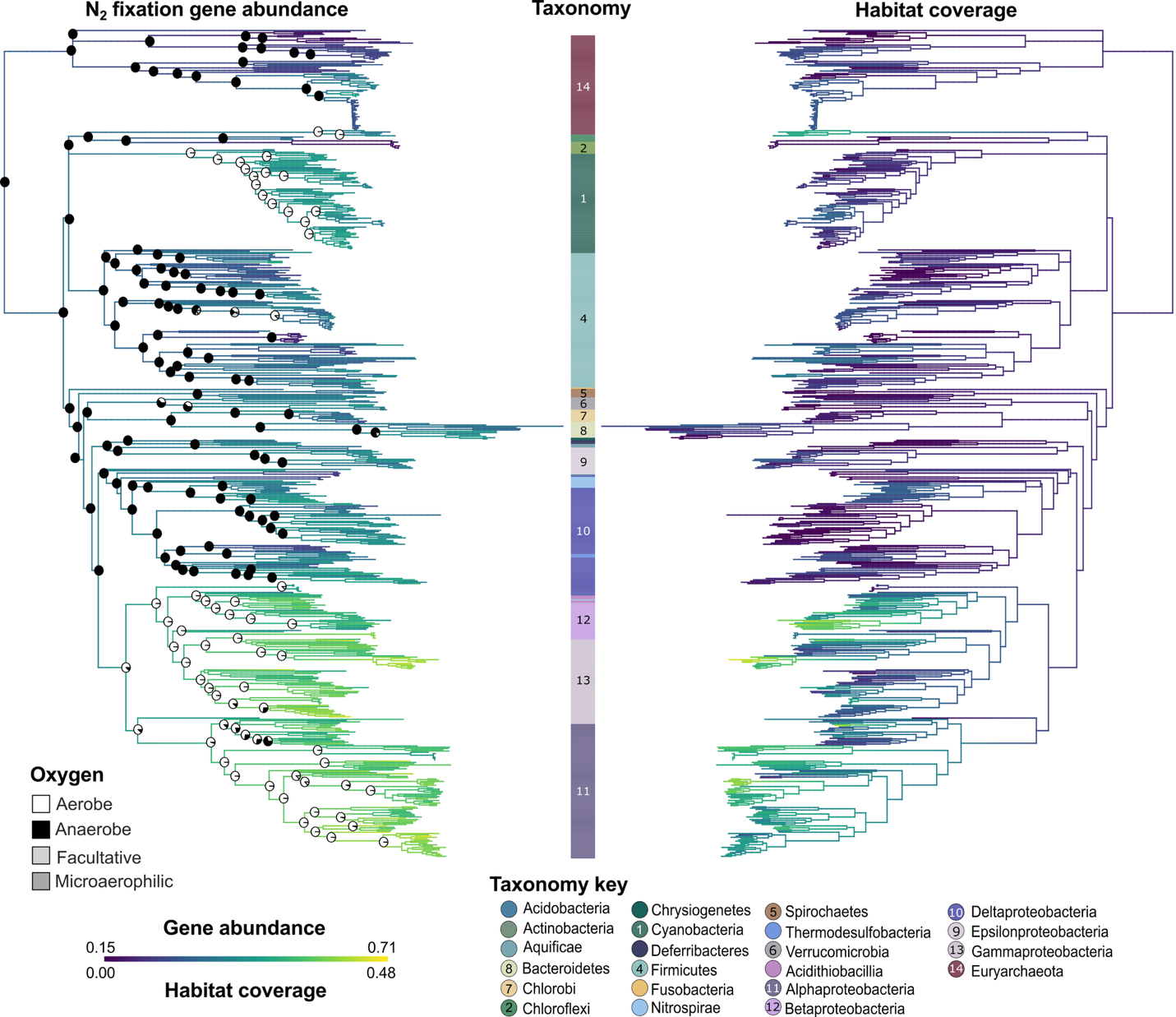

**Supplemental Figure 12.** Ancestral state reconstructions of N_2_ fixation gene abundance (right) habitat coverage (left) across the species phylogeny of nitrogenase-containing genomes. Darker branch colors represent a higher estimated gene abundance or broader habitat coverage. The *fastAnc* function in phytools was used for continuous variable reconstructions of metabolic and habitat diversity, which was mapped onto the tree using the *contMap* function [7]. On the gene abundance tree, reconstructed probabilities of ancestral oxygen preference are shown at selected nodes, with pie chart symbols representing the most probable states (aerobic, anaerobic, facultative, or microaerophilic). The *fitdiscrete* function in geiger [9] was used for discrete reconstructions of oxygen preference, which were mapped onto the phylogeny with the *simmap* function in phytools [7].
